# Impact of stimulation location relative to grey and white matter on single pulse electrical stimulation responses in the human brain

**DOI:** 10.1101/2021.10.07.463524

**Authors:** Angelique C. Paulk, Rina Zelmann, Britni Crocker, Alik S. Widge, Darin D. Dougherty, Emad N. Eskandar, Daniel S. Weisholtz, R. Mark Richardson, G. Rees Cosgrove, Ziv M. Williams, Sydney S. Cash

## Abstract

**Background:** Electrical neuromodulation is an increasingly common therapy for a wide variety of neuropsychiatric diseases. Unfortunately, therapeutic efficacy is inconsistent, possibly a result of our limited understanding of the mechanisms and the massive stimulation parameter space.

**Objective/Hypothesis:** To better understand the role different parameters play in inducing a response, we systematically examined single pulse-induced cortico-cortico evoked potentials (CCEP) as a function of stimulation amplitude, duration and location in the brain and relative to grey and white matter.

**Methods:** We measured voltage peak amplitudes and area under the curve of intracranially recorded stimulation responses as a function of distance from the stimulation site, pulse width, current injected, location relative to grey and white matter, and brain region stimulated (N=52, n=719 stimulation sites).

**Results:** Increasing stimulation pulse width increased response values near the stimulation location. Increasing stimulation amplitude (current) increased responses nonlinearly. Locally (<15 mm from the stimulation site), stimulation closer to the grey matter-white matter boundary induced larger responses. In contrast, for distant sites (>15 mm), white matter stimulation consistently produced larger responses than stimulation in or near grey matter. These relationships were different between cingulate, lateral prefrontal, and lateral temporal cortical stimulation.

**Conclusion:** These results demonstrate the importance of location and stimulation parameters in inducing a specific output and indicate that a stronger local response may require stimulation in the grey-white boundary while stimulation in the white matter may be needed for network activation, suggesting that stimulation location can be tailored for a specific outcome, key to informed neuromodulatory therapy.

**Highlights:**

1. Intracranial single pulse electrical stimulation (SPES) response increases with increased pulse duration mostly near the stimulation site
2. SPES response varies nonlinearly with injected current with an effect of distance from the stimulation site.
3. SPES near the grey-white boundary and 90° to the nearest cortical axis induces larger local responses, but white matter stimulation produces larger distant responses.
4. The relationship between SPES location and responses depends on brain region stimulated

## Introduction

Direct electrical stimulation (DES) of brain tissue can alleviate symptoms of neuropsychiatric diseases ranging from Parkinson’s to obsessive compulsive disorder (OCD) to epilepsy [1–11]. However, neurophysiological, behavioral and therapeutic effects of DES remain quite variable and unpredictable [12–17]. This variability in efficacy likely results from complex interactions amongst multiple factors, including stimulation location, pulse width and duration, injected current, location relative to grey and white matter, and stimulation frequency [14,18–33]. Attempts to better understand the mechanisms underlying DES neuromodulation are complicated by near infinite parameter possibilities as well as numerous hypotheses for why certain stimulation parameters are effective while others are not [2,6,8,13,34–38]. Adding to this challenge is that there are only a handful of studies that systematically examine the physiological effects of invasive brain stimulation in humans [24,28,39–46]. Not fully understanding the input-output relationships between DES and human cortical responses makes it difficult to predict which parameters, if adjusted properly, will optimize therapeutic effects.

A growing body of literature, however, has revealed consistent relationships between some stimulation parameters and neural responses across individuals [24,28,29]. This suggests that there are consistent, but only partially identified, maps for how the human brain responds to different stimulation parameters. This is best reflected in isolated single pulse electrical stimulation (SPES, via macro-electrodes) where local cortico-cortico evoked potentials (CCEPs) usually show consistent waveforms divisible into N1 and N2 voltage peaks [29,47–53]. In addition, SPES have been extensively used to map brain connectivity including the effect of pathologies, such as epilepsy [52,54]. Various parameters affect the CCEP size and shape [29,47–53].

Despite the importance of SPES, there is little work systematically exploring the relationship between input parameters and physiological output. For instance, there is little information on the impact of stimulus duration on CCEP. Pulse width, or duration, has been shown to have an effect in mouse models as well as a possible clinical effects in Parkinson’s Disease [32,33,39,55]. However, with SPES, pulse width ranges from 0.3 to 3000 ms have been reported as well as both monophasic and biphasic stimulation with few consistencies across studies in the field [44,51,52,56]. Therefore, there remains a need to use the same stimulation parameters and approach to identify the relationships between responses and SPES pulse width duration parameters. Our expectation is that increasing duration will induce larger and more widespread stimulation responses.

Similarly, little is known about how the amplitude of injected current sculpts the shape of the voltage response. With trains of stimulation (which include multiple single pulses), injected current has been shown to have a linear relationship with behavioral and physiological responses [24,27]. Therefore, we hypothesize that there would be a linear relationship between SPES responses and current amplitude.

Stimulation location effects on responses is expected to be more complex, as DES location can be relative to the ‘microarchitecture’ (e.g., cortical layer or nearness to the closest white matter tract) or ‘macroarchitecture’ (e.g., location relative to other brain regions or white matter bundles reaching those regions). Stimulation location, not surprisingly, has a substantial impact on both behavioral and physiological responses with differing clinical outcomes [14,28,42,51,57–60]. The most therapeutically effective DES, whether inducing very focal or more widespread neural responses or engagement of the network to induce different states, is thought to generally engage white matter tracts [29,60], which has been supported by a recent study relating stimulation effects to white matter proximity [28]. Indeed, some experiments and modeling studies of electrode location, spacing, and orientation relative to subcortical regions and white matter tracts have suggested it is possible to map, and plan, DBS approaches to subcortical stimulation that predict optimal therapeutic effects [6,39,61,62]. But parallel information for cortical structures is lacking.

Further, there is an effect of distance from the stimulation site that must be taken into account. Responses close to the site of stimulation are quite different from ones further away and their dependence on parameters may differ as well. For example, local stimulation responses appear to reflect functional connectivity while distant stimulation responses may better reflect structural (white matter tract) connectivity [29].

Based on this existing literature, we hypothesized that: 1) increasing waveform single duration (pulse width) engages more of the distant brain network; 2) SPES responses linearly vary with injected current locally and distally; 3) location of the bipolar pair of stimulation electrodes in grey matter is lower than responses to stimulation closer to white matter (along a linear curve) with a linear effect of orientation on the responses; 4) these relationships depend on the region stimulated. To test these hypotheses, we sampled the neural responses to intracranial SPES via macroelectrodes across a population of patients with intractable epilepsy (N=52) implanted with intracranial leads in the process of monitoring for seizure foci across 719 stimulation sites. Focusing on the bipolar, charge-balanced stimulation between neighboring macroelectrodes used in numerous studies [24,49,51,52,63–65], we examined neural responses to stimulation across >6500 bipolar recording sites.

## Materials and Methods

### Human Participants and Recordings

We recorded intracranial neural activity from participants with intractable epilepsy undergoing invasive monitoring. A subset of the data from the single pulse electrical stimulation data were used in a previous publication (N=11) [29]. Most participants were implanted with multi-lead depth electrodes (aka sEEG) with a smaller subset implanted with either grid or strip electrodes (a.k.a. ECoG) to locate epileptogenic tissue in relation to essential cortex. Data using stimulation in the the grid or strip electrodes were not included in the analyses here. Depth electrodes (Ad-tech Medical, Racine WI, USA, or PMT, Chanhassen, MN, USA) with diameters 0.8–1.27 mm and 4-16 platinum/iridium-contacts 1-2.4 mm long with inter-contact spacing ranging from 4-10 mm (median 5 mm) were stereotactically placed in locations deemed necessary for seizure localization by a multidisciplinary clinical team independent of this research. Following implant, the preoperative T1-weighted MRI was aligned with a postoperative CT using volumetric image coregistration procedures and FreeSurfer scripts ([66– 69]; http://surfer.nmr.mgh.harvard.edu). Electrode coordinates were manually determined from the CT in the patients’ native space [69] and mapped using an electrode labeling algorithm (ELA; [70,71]) that registered each contact to a standardized cortical map [72].

Participants received their normal antiepileptic medications prior to stimulation testing to minimize the risk of seizure. Recordings used a Blackrock system with a sampling rate of 2 kHz (Blackrock Microsystems, Salt Lake City, UT, USA). Depth recordings were referenced to an EEG electrode placed on skin (C2 vertebra or Cz), a chest EEG lead contact, or via an internal ground.

### Ethics statement

All patients voluntarily participated after fully informed consent as monitored by the Partners Institutional Review Board covering Brigham and Women’s Hospital (BWH) and Massachusetts General Hospital (MGH). Participants were informed that participation in the stimulation tests would not alter their clinical treatment in any way, and that they could withdraw at any time without jeopardizing their clinical care.

### Neural stimulation

Stimulation was delivered with a CereStim stimulator (Blackrock Microsystems, Salt Lake City, UT) using single pulse electrical stimulation (SPES). Current injection and return paths used neighboring contacts in a bipolar configuration [42]. Stimulation was controlled via a custom Cerestim API via MATLAB or a custom C++ code (https://github.com/Center-For-Neurotechnology/CereLAB). Waveforms of two different durations were used: 1) 233 µs duration: 90 µs charge-balanced biphasic symmetrical pulses with an interphase interval of 53 µsec with between 5 and 100 trials with a median of 20 trials per stimulation site (mean number of trials: 18.9±12.21) [29,64], and 2) 1053 µs (∼1 msec) duration: 500 µs charge-balanced biphasic symmetrical pulses with an interphase interval of 53 µsec with between 10 and 26 trials per stimulation site with a median of 10 trials per site (mean number of trials: 10.4±3.49) [52,73– 75]. The interval at 53 ms was required as a hardware-limited minimum interval between square pulses with the CereStim stimulator. Multiple current amplitudes were applied with the short duration (233 µsec) bipolar stimulation at the following steps: 0.5 mA to 10 mA at 0.5 mA steps with a minimum of 10 trials per stimulation site (mean number of trials: 14.7±5.64). The two durations were tested in ten participants and current amplitudes were tested in eleven participants. A trained electroencephalographer examined ongoing recordings for epileptiform activity and asked participants if they experienced any sensations. The participants were awake and were aware that they were being stimulated but were blind to the stimulation timing and parameters.

Electrode locations for stimulation were chosen to avoid known areas of seizure onsets as judged by the participants’ clinicians. We also avoided stimulation in sites which were in or near the corpus callosum as well as sites which were too medial in the cingulate near the meninges as it could cause discomfort.

### Stimulation location and electrode measures

For identification of electrode location relative to grey and white matter, we measured the orthogonal Euclidean distance from the center of each bipolar pair of electrodes to the nearest reconstructed vertex of the pial and white matter surfaces generated from FreeSurfer tools following colocalization [66–69]. As the depth electrode can curve during implantation, the Euclidean distance between contacts can change. Therefore, bipolar stimulation pair distances between each contact were re-calculated using Euclidean measures to compensate for the depth electrode curving or bending slightly when implanted. The orientation (angle) of the bipolar pair of electrodes relative to the cortical column was calculated by first detecting the nearest grey-white and outer surface (pial) points to the center of the bipolar stimulation pair. A line perpendicular to the tangent was calculated from the base (edge of layer 6) and the outer axis of the cortex (pial, outer edge of layer 1) which formed the cortical column pole. Then, the angle between the two poles (the cortical column and the bipolar pair) was calculated using the dot product of the two line segments. We also categorized stimulation sites as in the grey matter, subcortical regions, white matter, and pial surface by identifying the colocalized stimulation location relative to the grey matter surface volume, the white matter surface volume, and the reconstructed subcortical volumes in the participants’ native space [66,67,69]. The classification of the stimulation sites relative to the surfaces was done using the MATLAB inpolyhedron function ([76], MATLAB 2020b). In addition, we used the MATLAB functions alphaShape and inshape to identify electrodes on the pial surface but not within the cortex.

### Data Analysis

Data analysis was performed using custom analysis code in MATLAB and Fieldtrip (http://www.ru.nl/neuroimaging/fieldtrip; [77]). Channels with excessive line noise or without clear neural signal (determined by visual inspection) were removed from the analysis. In addition, we removed pathological channels with interictal epileptiform discharges (IEDs) using an automatic IED detection algorithm ([78], version v21, default settings except -h at 60; http://isarg.fel.cvut.cz). We removed from analyses channels which had detected IEDs greater than 6.5 IEDs/minute (which is above the algorithm’s false positive rate). These checks resulted in mean=8.25 ± 11.411 channels removed per participant. The remaining electrodes were subsequently demeaned and bipolar re-referenced relative to nearest neighbors to account for volume conduction [79,80]. Stimulation artifact was removed in the 20 ms around the onset of the pulse for the single pulses using a Tukey-windowed median filter [81].

The CCEP, a type of event-related potential (ERP), was analyzed by extracting epochs from 1000 ms before stimulation onset to 3000 ms after offset. Voltages were normalized by z-scoring the voltage values using the baseline data segment per trial. The number of trials per condition ranged from 5 to 100 trials, median of 20 trials across conditions (conditions include either duration, current amplitude, or location). To remove trials that had spurious voltage deflections that could be due to acute spurious external events such as referential noise and were not just a voltage response relative to baseline, we rejected trials where the voltage exceeded 5 standard deviations from the average response of all trials per site (not the baseline) per channel and condition. The average number of trials removed across participants was 2.3±1.17. The average ERP across the remaining trials per condition was then calculated and the following metrics were measured for the average ERP: i) normalized overall peak amplitude (the peak amplitude during the full second after stimulation), ii) area under the curve (sum of the absolute values of zscored data for 1 sec after stimulation offset), iii) absolute value of the N1 peak (between 10 and 50 ms after stimulation), iv) and absolute value of the N2 peak (50-400 ms post-stimulation) [24,49,52].

Based on an examination of the voltage amplitudes relative to distance, we used a data-driven cutoff where neural data were subdivided into local responses (< 15 mm from the stimulation site) and distant (> 15 mm from the stimulation site), to separate the waveform characteristics nearest to the stimulation site from propagated activity.

### Statistical analysis

We tested comparisons across brain regions and stimulus parameters with the Kruskal– Wallis test for non-equivalence of multiple medians followed by *post hoc* Tukey-Kramer tests to determine statistically separable groups. A significant post stimulation response was identified as being significantly above or below baseline activity using Wilcoxon rank sum comparisons. Multi-way ANOVA was used to examine effects of different parameters on the neural responses to determine whether brain region and stimulation parameters have stronger effects on neural activity. We corrected for multiple comparisons at a target p-value (0.05) with a Bonferroni correction. For spatially resolved analyses, we applied a false discovery rate (FDR) correction to the p-value at each spatial step, identifying significance as that point where the FDR-corrected value was lower than the Bonferroni-corrected target (as there were four different measures).

To detail the relationships between stimulation parameters (duration, current, location) and neural responses, we performed Pearson’s linear correlation between the parameters and the responses (whether voltage measures). We also performed a series of model fits using generalized linear regression models (GLM), using the functions ‘fitglm’ (assuming a normal distribution and an identity linkage to generate the first through tenth order models) and the function ‘predict’ to produce the fitted curves to the mean responses per stimulation site to the grey/white matter boundary location. To identify the relationship between the distance to the grey-white boundary and neural responses (AUC, overall peak, N1 peak, and N2 peak values) relative to the distance to the stimulation site, we fit the models to different distance thresholds distinguishing ‘local’ versus ‘distant’ recording sites at different distances from the stimulation site (e.g. 15-50 mm). We identified the best model as that one which had the lowest Aikake Information Criterion (AIC). We then used F-test to test whether the model was a better fit to the data compared to a model which only included a constant term (indicating there was no relationship between the variables). Every fitted line was to the data points which were averaged per stimulation site per condition, such that the recording sites were binned based on stimulation location and distance relative to the stimulation location.

## RESULTS

We performed direct electrical stimulation (DES) while recording from intracranial leads in patients with intractable epilepsy undergoing clinically indicated neural monitoring to delineate the seizure focus (N=52; median age= 37, ranging from 18 to 67; 30 women). Only data from implanted stereoEEG (sEEG) depth electrodes were examined in this data set (**Fig. 1**). Brain region coverage of the total of 719 individual stimulation locations across participants was relatively widespread, with a concentration of stimulation sites in the medial temporal lobe and lateral prefrontal lobes; **Fig. 1A; Supplemental Table 1**). We examined neural responses to DES at different stimulation sites across >6500 bipolar re-referenced recording sites (termed channels; **Fig. 1B**). We subdivided the stimulation sites into six main regions: lateral prefrontal lobes (includes the lateral prefrontal cortex and insula), cingulate cortex, lateral temporal lobe (including the lateral middle, superior, and inferior gyri), parietal lobe, subcortical areas (which includes the hippocampus, amygdala, caudate, and putamen), and occipital lobe (**Fig. 1A; Supplemental Fig. 1**). The SPES responses were measured as average absolute peak (the overall peak), area under the curve (AUC), N1 peak, and N2 peak (**Fig. 1C**).

**Figure 1.**
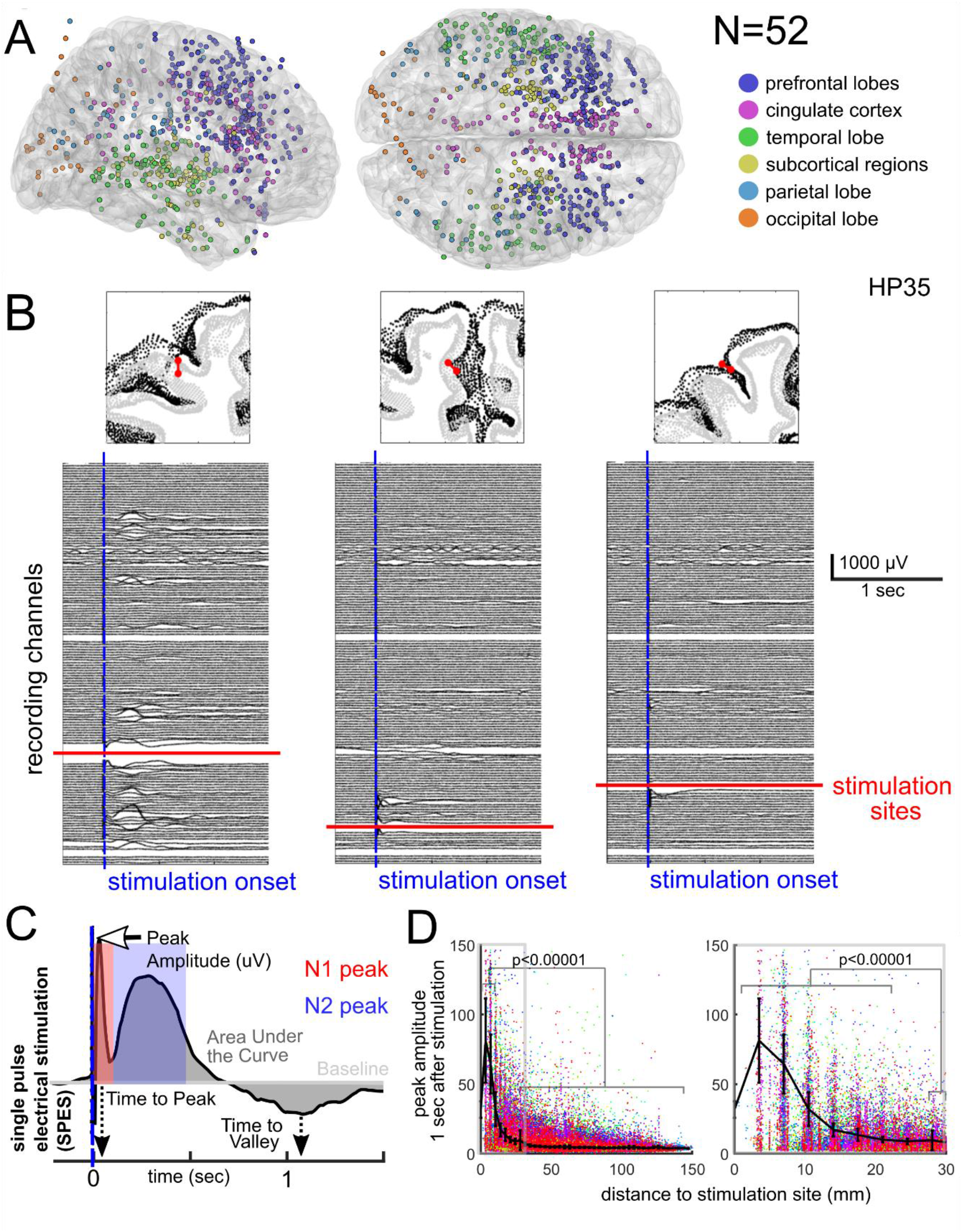
Single pulse electrical stimulation (SPES) responses vary nonlinearly with stimulation distance. **A**. Locations of all SPES stimulation locations (n=719) overlaid on the colin27 brain and divided into six different brain region categories, N=52. Electrode locations are only approximate as shown here as the locations were not morphed to a common brain. **B**. Top: locations of the stimulating electrodes (red dots with a line) in coronal slices, with black dots indicating the pial surface and grey-white boundary. Left and right columns are lateral prefrontal cortex and middle is cingulate. Bottom: Average stimulation responses after bipolar re-referencing (20 trials) for each location in the same participant (HP35). Red lines indicate the relative location of the stimulating electrodes to the recording electrodes (in black lines). Blue lines indicate stimulation onset. **C**. Example of the measurements of the cortico-cortico evoked potential (CCEP) following SPES. **D**. Distance dependence for responses for the overall peak responses during the 1 second after stimulation, with color coding representing different participants (N=52, n=719 stimulation sites). Left is the entire range up to 150 mm while the right focuses on 0-30 mm. Error bars are mean and S.D. of the distributions across participants; p-value based on Kruskal-Wallis test followed by a *post hoc* Tukey Kramer test.

One attribute that must be considered is the fundamental difference in local versus distant responses [29]. In particular, there can be substantial differences in response spatial extent from stimulating electrodes at different sites (**Fig. 1B; Supplemental Fig. 2**; HP35). Therefore, we performed a preliminary analysis of the distance between the recording and the stimulating electrodes across the data set. We found a significant difference in the responses between recording sites <21 mm and sites >35 mm away from the stimulation site with an inflection point around 15 mm, whether the contacts were along the same depth electrode or not (p<0.00001; Kruskal-Wallis test; overall peak: Chi-Sq=1800.32; N=52; **Fig. 1D**; **Supplemental Fig. 3**,**4**). Hypothesizing that local and distant responses could indicate different rules for stimulation parameters, as suggested in past studies, [28,29,49,65,82,83], we used this data-driven boundary to subdivide the responses into local and distant recording sites to address four main hypotheses: 1) increasing waveform duration engages more of the distant brain network; 2) SPES responses linearly vary with injected current locally and distally; 3) electrode orientation and location with regard to white and grey matter varies linearly with responses to the stimulation locally and distally; 4) these relationships depend on the brain region stimulated with similar relationships locally and distally. We also examined these questions relative to a continuous measure of distance.

### Stimulation Duration has a Stronger Effect on Local Sites

We applied SPES at two different durations at multiple stimulation sites in a subset of participants: 233 µsec and 1 ms (N=10, 9.3±5.14 sites per participant). With the increase of duration, local (<15 mm) and distant (>15 mm) responses were significantly increased as measured by the overall peak, the AUC, the N1 peak, and the N2 peak (p<0.00005; Wilcoxon rank sum test; N=10; n=102 stimulation sites; n=1273 recording sites; **Fig. 2A-D**). Examining multiple points away from the stimulation site, the difference between the shorter and longer durations diminished, to the point that there no difference between durations at distant sites (**Supplemental Fig. 3B**). This was likely due to the fact that there was little to no response past a certain distance from the stimulation site [29].

**Figure 2.**
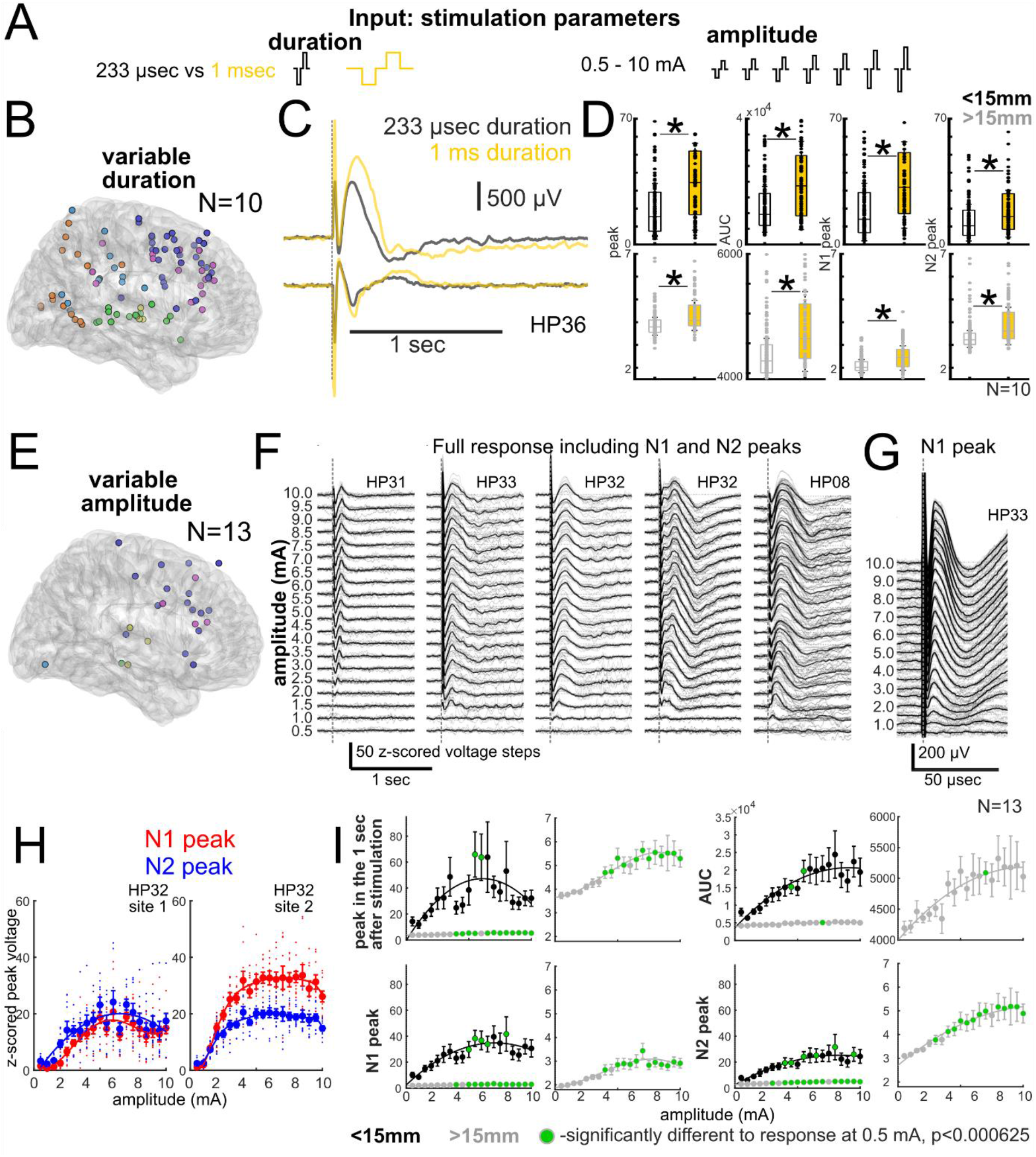
Responses vary nonlinearly with current, stimulation duration and recording site distance. **A**. Schematic of amplitude steps and stimulation durations. **B**. Locations of all SPES stimulation locations where we tested both durations (233 µsec versus 1 ms) mapped to a single brain (N=10; 102 stimulation sites), with six different brain regions colored as above. **C**. Example of stimulation responses to two different durations (233 µsec, black, 1 ms, yellow) recorded at two neighboring sites (participant HP36). **D**. Overall peak, AUC, N1 peak, and N2 peak responses for the two durations for local sites (black lines, upper plots) and distant sites (grey, bottom plots). * indicates p<0.00005, Wilcoxon rank sum. **E**. Locations of all SPES stimulation locations where we tested multiple current steps mapped to a single brain (N=13; 25 stimulation sites). **F**. Average (black line) and individual trial stimulation responses (grey lines) at neighboring electrodes for 20 amplitude steps (0.5-10 mA, with 0.5 mA steps, median 10 trials per step) for five sites in the lateral prefrontal lobe and four participants (designations HPXX), largely showing N2 responses. **G**. Higher time resolution of responses to the 20 amplitude steps for one site in the lateral prefrontal lobe (participant HP32) showing the N1 peak more clearly. For **F** and **G**, grey lines are per trial and black lines are average responses. **H**. N1 (red) and N2 (blue) peaks at different amplitude steps at neighboring contacts for two lateral prefrontal lobe stimulation sites, one in each hemisphere, in the same participant. Dots are per trial, error bars indicate mean and standard error per current step. Continuous curves are generalized linear model (GLM) second order polynomials. **I**. Overall peak (peak zscored voltage in the full 1 second after stimulation), AUC, N1 peak (peak in the 50 ms after stimulation), and N2 peak responses (peak in the 50 ms-250 ms after stimulation) for local (<15mm, black) and distant (>15mm, grey) recording sites. For each measure (overall peak, AUC, N1, N2), the left plot includes the <15 mm and >15 mm recording sites and the right plot includes an expanded view of the >15 mm response. Every voltage value per trial is z-scored (and therefore zero-mean corrected) relative to the baseline value. Error bars indicate mean and standard error per current step; green dots are current levels significantly different from the responses at 0.5 mA for the same response measure and distance, p<0.000062 (multiple comparisons corrected).

### CCEP Response Features are Nonlinearly related to Increasing Injected Current

To test the hypothesis that the size of the evoked potentials tracks linearly with increasing stimulation current, we examined the relationship between current amplitudes and responses at a set pulse with duration (233 µsec), stepping from 0.5 mA to 10 mA in 0.5 mA increments (N=13 participants, 1.6±0.89 DES sites per participant, median 10 trials per current step; **Fig. 2A, E-G**). First focusing on the immediate neighboring recording channels, we found increasing current induced larger voltage responses for both the N2 peak (**Fig. 2F**) and the N1 peak (**Fig. 2G**), though the response-current relationships were not linear, but appeared more to be an s-shaped curve (**Fig. 2H**). A threshold around 1-2 mA had to be reached to induce a clear CCEP. Between ∼0.5 and ∼5 mA, the increase in response with injected current was linear. Beyond ∼5 mA, however, the responses plateaued or even decreased (**Fig. 2H**).

Moving beyond single neighboring contacts to all contacts, we found a nonlinear DES current-CCEP response relationship for overall peak, AUC, N1 peak, and N2 peak (**Fig. 2I**). To emphasize that the relationship between response and current for the entire current range was nonlinear, the Pearson’s linear correlation between the current and response was low (mean 0.14±0.04). Interestingly, the responses peaked in local (<15 mm) recording sites, where we found the responses to the middle current ranges (5-8 mA) were significantly different relative to the lowest current level (0.5 mA), but not the highest current responses (Wilcoxon rank sum test; p<0.000062; **Fig. 2I**). In contrast, distant sites had significant responses (above the response at 0.5 mA) around 3-4 mA (for N1 and N2 peaks) to 5-7 mA (for AUC and overall peaks) which remained high and plateaued for the peak measures (Wilcoxon rank sum test relative to responses at 0.5 mA; p<0.000062; **Fig. 2I**). These nonlinear relationships was further demonstrated by the fact that, in local sites (<15 mm), a second order polynomial (using a GLM) better fit the relationship between increasing current and the overall peak, AUC, N1 peak, and N2 peak (**Fig. 2I**, black line) while third order polynomial S-shaped curve best described the current-response profiles for distant recording sites (**Fig. 2l**, grey line) based on the minimum AIC following the GLM fit; n=25 stimulation sites; n= 1797 recording sites).

At finer spatial steps away from the stimulation electrode at 4 mm steps from 4 to 100 mm, we found increasing current amplitude had a stronger effect in local recording sites, with significant differences between amplitudes <39 mm away from the stimulating electrode (p<0.00005; Kruskal-Wallis multiple comparisons test; maximum distance with a significant difference between amplitudes: overall peak: max dist.=36 mm; AUC: max dist.=36 mm; N1 peak: max dist.=24 mm; N2 peak: max dist.=39 mm; **Supplemental Figure 3**). These results once again demonstrate that a different set of stimulation parameter-response rules govern local versus distant stimulation responses though, contrary to our original hypothesis, responses vary nonlinearly with stimulation current amplitude, especially locally.

### Stimulation Responses Vary with Location in Cortex and White Matter

We tested the relationship between responses and DES location relative to grey and white matter. Controlling amplitude and duration parameters while focusing on location, we examined responses at 7 mA at the shorter pulse duration of 233 µsec for 719 independent stimulation sites across 52 participants (14.5±10.81 DES sites per participant; 6772 recording sites; **Fig. 3A**; see **Methods**). We chose these values since this combination of parameters consistently produced a robust response past 6 mA, even with 233 µsec pulse duration (**Fig. 2**). Choosing different stimulation sites along the depth electrode that were in different portions of the cortical column and white matter, we found larger N2 peak responses in the white matter and at the grey-white boundary compared to stimulation in the grey matter alone (example participant in **Fig. 3A**).

**Figure 3.**
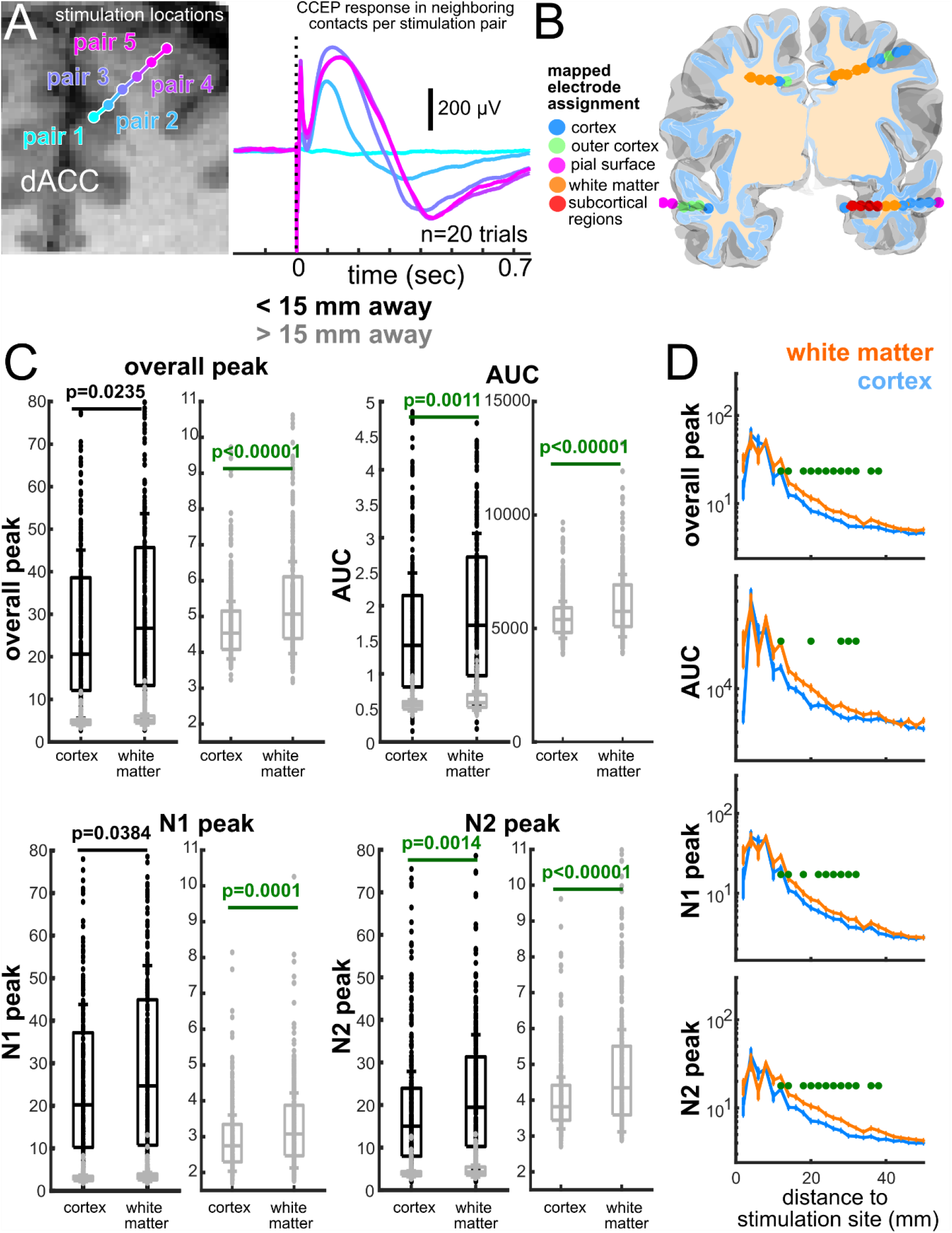
Responses depend on stimulation location along grey-white axis. **A**. Stimulation locations and responses in example participant (HP35). Evoked potentials averaged across 20 trials. **B**. Categorization of electrode localization, with an illustration of the cortical and white matter volumes relative to implanted leads and example categorization of implanted leads. **C**. Effect of stimulation site on peak neural responses (upper left panel), area under the curve (AUC; upper right panel); N1 peak (lower left panel) or N2 peak (lower right panel). Recording sites are separated into local (<15mm away from the stimulation site; white bars with black outlines, top row of figures) or distant sites (recording sites >15mm away from the stimulation site, grey outlined bars, and expanded to the right). For each measure (overall peak, AUC, N1, N2), the left plot includes the local and distant sites while the plot to the right an expanded view of distant responses. Each dot is per stimulation site (not per recording site). Green p-values are significantly different after correcting for multiple comparisons. **D**. Effect of changing stimulation location relative to the cortex and white matter on responses and distance relative to the stimulation. The difference is significant between the different volumes at stimulation-recording electrode distance steps between ∼15 and 50 mm (green dots, p<0.00005). All stimulation is at 7mA and 233 µsec; N=52; p-values are from Wilcoxon rank-sum test.

To quantify this dependence on location, we used two approaches, one categorical and one a continuous distance measure which also allowed us to account for the wide variety of electrode placement as present in the data set. In the first, we categorized stimulation sites as being on or near the pial surface, in the outer cortex (with one contact of the bipolar depth at or near the outer edge of the cortex, though this could also include being within a sulcus), fully in cortex, in white matter, and in subcortical regions (amygdala, hippocampus, caudate, and putamen) in the participants’ native space (see **Methods; Fig. 3B; Supplemental Figure 1, 2**). For the second approach, we measured the Euclidean distance of the stimulation site from the grey-white boundary. Both approaches demonstrated that responses increased with proximity to white matter overall.

With the categorical method, at local recording sites (<15 mm), we found the AUC and N2 peak were significantly higher with white matter stimulation versus cortical stimulation (p<0.0014; Wilcoxon rank sum test), but not the overall peak and N1 peak responses after correcting for multiple comparisons (p>0.0063; Wilcoxon rank sum test; **Fig. 3C**; **Supplemental Figure 5**). In contrast, responses at distant recording sites (>15 mm) were significantly higher when stimulation was in white matter and subcortical regions compared to cortex for all measures (p<0.0001; Kruskal-Wallis Multiple comparisons test; post hoc Tukey-Kramer test; **Fig. 3C; Supplemental Figure 5**). The lowest stimulation responses occurred when the bipolar pair was in the outer cortex, which could include the depth contacts being within a sulcus and just at the pial surface (**Fig. 1B, 3A; Supplemental Figure 1, 2; Supplemental Figure 5**). For all locations, stimulation in the white matter compared to cortex induced an average of 13.1±2.1% higher responses across voltage measures. For distant (>15 mm) sites only, this was a 31.1 ±1.6% increase while only an 4.9±0.5% increase for local sites (Local responses: Overall peak: cortex: 26.8±1.79; white matter: 27.6±1.62; AUC: cortex: 1.8 × 10^4^±0.11 × 10^4^; white matter: 1.7 × 10^4^±0.10 × 10^4^; N1 peak: cortex: 25.3±0.89; white matter: 26.2±1.54; N2 peak: cortex: 17.5±1.05; white matter: 15.9±0.93; Distant responses: Overall peak: cortex: 3.0±0.18; white matter: 3.8±0.22; AUC: cortex: 2.2 × 10^3^±0.01 × 10^3^; white matter: 2.7 × 10^3^±0.02 × 10^3^; N1 peak: cortex: 2.7±0.16; white matter: 3.5±0.20; N2 peak: cortex: 2.1±0.13; white matter: 3.2±0.18; **Fig. 3C; Supplemental Fig. 5**).

### Stimulus Location has Largest Effects at Intermediate Distances

In the foregoing results, stimulation in white matter leads to larger responses locally (<15 mm) but only for N2 peak and AUC, while at a distance (>15 mm) this is true for all measures. Does white matter stimulation lead to larger responses at any given long distance? To answer this question and to understand if changing the stimulation location also changes the degree to which activity spreads across the brain, we compared evoked responses at a given distance from the site of stimulation (**Fig. 3D; Supplemental Fig. 6**). Between 15 and 40 mm from the stimulation site, there were significantly larger responses when stimulation was in the white matter compared to cortical stimulation (p<0.000063; Wilcoxon rank-sum test for comparing only white matter versus cortex at different distances; **Fig. 3D**). For recording sites less than 15 mm and more than 40 mm away from the stimulation site, there was no statistically verifiable difference between stimulation in grey vs. white matter. Expanding the response comparisons to include the pial surface, outer cortex, cortex, white matter, and subcortical categorizations, responses at a distance between 15 and 40 mm from the electrode were significantly larger with white matter stimulation compared to other structures (pial, outer cortex, and cortex) stimulation (p<0.00005; Kruskal-Wallis test when comparing multiple volumes; **Fig. 3D; Supplemental Fig. 6**).

### Stimulation Responses Varies with Distance to the Grey-White Matter Boundary

As discussed above, in addition to the categorical approach which produces an ordinal variable, we used an interval variable measuring the Euclidean distance between the center of the stimulation bipolar pair and the grey/white boundary (**Fig. 4A; Supplemental Figure 7**). We adopted a convention using an axis spanning the grey-white boundary wherein negative distance values indicate cortex while positive values are in the white matter and the grey-white junction is 0 (**Fig. 4; Supplemental Figure 7**). Corroborating the results using the categorical approach, we found a significant positive correlation between the proportion of channels with above-threshold stimulation responses (average response 5 STD above the mean baseline activity before stimulation) and stimulation location relative to white matter (Pearson’s correlation, rho=0.16, p=0.00002; **Fig. 4B**).

**Figure 4.**
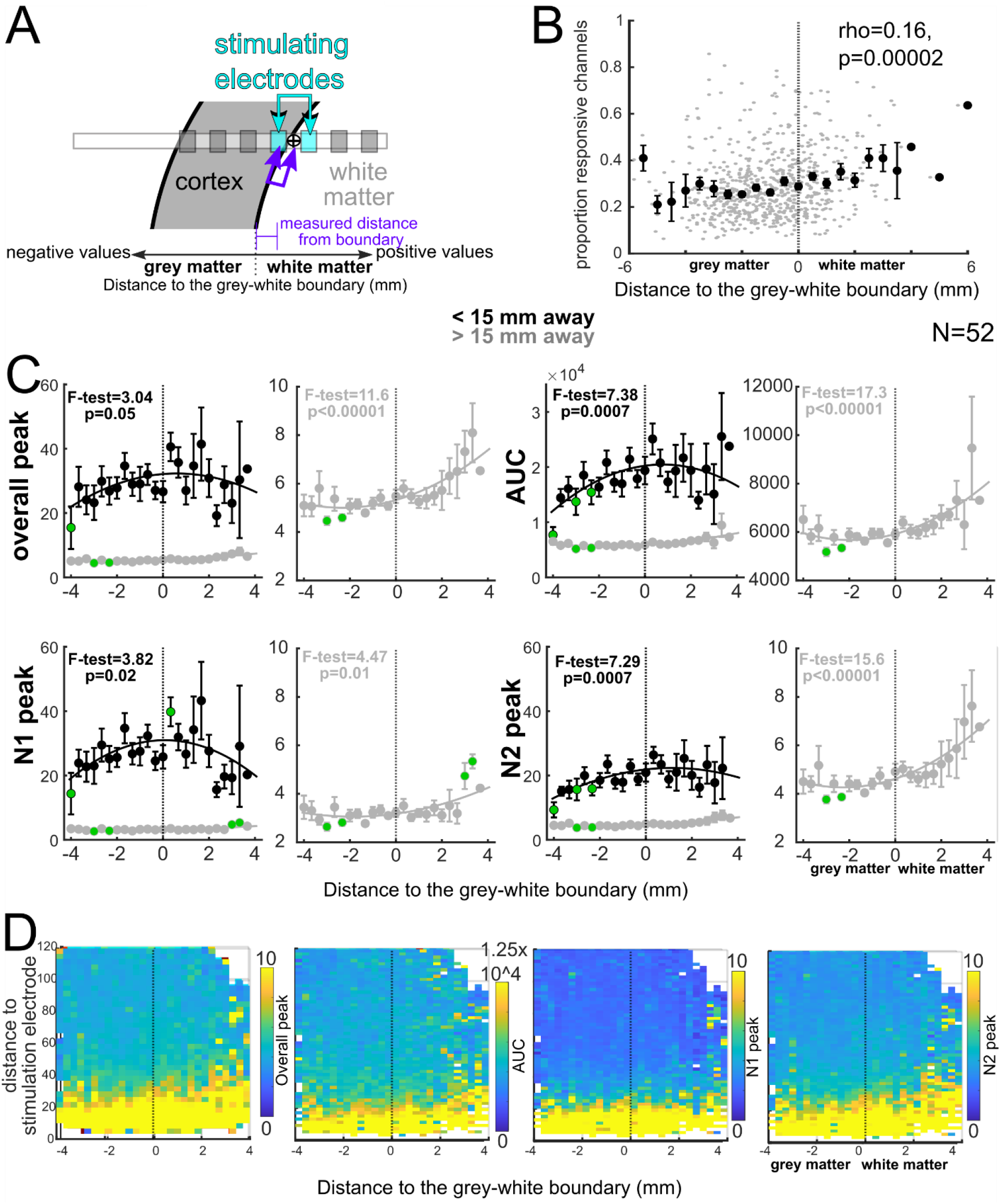
Responses are non-linearly related to stimulation location along the grey-white axis. **A**. Schematic of localization measurements. **B**. Proportion of all channels per stimulation site and participant with above-threshold responses. **C**. Local (<15 mm, black lines and curves, left) and distant (>15 mm, grey lines and curves, expanded to the right) peak in the 1 second after stimulation (top left), AUC (bottom left), N1 peak (top right), and N2 peak (bottom right) relative to the distance to the grey-white matter boundary. For each measure (overall peak, AUC, N1, N2), the left plot includes the <15 mm and >15 mm recording sites and the right plot includes an expanded view of the >15 mm response. Data is from all contacts, not just the above-threshold responses. Green dots-responses at different distances to the grey-white boundary significantly different to responses at -4 mm, p<0.000062 (Wilcoxon rank sum test; multiple comparisons corrected). F-statistics and p-values for the model fits as compared to a model with a constant term for the local (black print) and distant (grey print) sites. **D**. Binned distance from stimulation site versus distance to the grey-white boundary for the average overall peak (peak in the 1 second following stimulation), AUC, N1 peak, and N2 peak responses across brain regions. Colorbar to the right of each plot indicates the measure and scale.

Examining the neural responses across the grey-white boundary for all local (<15 mm) recording sites, we found, contrary to our original hypothesis,a nonlinear relationship between this distance to the grey-white boundary and responses wherein the Pearson’s linear correlation values between responses and distance to the grey-white boundary were low (mean rho for all voltage measures=0.08±0.022; N=52). Instead, after fitting data to multiple models using a GLM fit and choosing the model order with the lowest AIC value, we found a second order polynomial described by an inverted u-shaped curve best fit the relationships between grey-white boundary axis and the AUC, overall peaks, N1 peaks, and N2 peaks. However, only the fits for the N2 and AUC responses relative to the grey-white boundary were significantly different to a constant term model (p=0.0007; **Fig. 4C**), indicating again that the stimulation location relative to white matter seems to most affect local N2 and AUC responses but not the local N1 or overall peak responses (**Fig. 3C**).

Distant (>15 mm) responses were also best fit by a second order polynomial using GLM though the curve resembled an exponential term for all measures, resulting in a substantial increase as stimulation location ‘moved’ into white matter (**Fig. 4C**; N=52). This curve contributed to the higher average Pearson’s linear correlation values for distant sites compared to local sites (mean rho for all voltage measures=0.15±0.037; N=52). Only the N1 response curve was not significantly different to a constant term model (F-test=3.82; p=0.01) while all other metrics followed a second order increasing polynomial fit and the relationship was significantly different from a constant term model (F-test; p<0.00001; **Fig. 4C**).

### Response Spread Increases with White Matter Stimulation

As was true with our previous analyses (**Fig. 4D**), responses at a distance were larger with white matter stimulation compared to cortical stimulation with the curves generally resembling an exponential or second order polynomial relationship. However, we found the relationship shifted from a second order polynomial (local) to a linear relationship as we included recording electrodes further away from the stimulating electrodes which was most prominent for the AUC measure (**Supplemental Figure 8**). Taking the recording electrodes within a range of distances (±5mm) at multiple steps away from the stimulating electrode, we found the models which best fit the relationship between responses and distance to the grey-white boundary could be linear, quadratic, or a second order polynomial (p<0.0001; fits shown by minimizing AIC and an F-test comparing the fitted models to a model only containing a constant term; **Supplemental Figure 8**). Therefore, in testing our original hypothesis that the relationship between response and distance to the white matter is linear, we once again found this is a non-linear relationship.

### Brain Region Impact on Stimulation Response Sensitivity to Grey-White Location

In the preceding results, we tested our hypotheses pooling all data regardless of brain region. Increasing literature, however, as well as inferences from known cytoarchitectural and connectivity differences would suggest that the rules for stimulation differ in different brain regions [24,28,51,60,84–87]. We explored this question by specifically comparing responses when stimulation was in lateral prefrontal lobe, cingulate, and lateral temporal lobe (as we had the most coverage of these regions; **Fig. 1A, Supplemental Figure 1**). We found that local responses were not significantly different between cortex and white matter stimulation in the cingulate or lateral temporal lobe for all measures (cingulate comparisons: p>0.64; temporal lobe comparisons: p>0.008; Wilcoxon rank-sum test), but were significant for the AUC and N2 peak responses with stimulation in the lateral prefrontal lobe (p<0.0024; multiple comparisons; Wilcoxon rank-sum test; **Supplemental Figure 9**). For distant sites, the overall peak and N2 peak responses were significantly higher with stimulation in the white matter compared to cortex for all three brain regions (p<0.004; Wilcoxon rank-sum test). Yet, the distant N1 peak response was not significantly different after correcting for multiple comparisons(p>0.01; Wilcoxon rank-sum test). AUC responses were only significantly higher with white matter near the lateral temporal lobe and cingulate (p<0.003; Wilcoxon rank-sum test; **Supplemental Figure 9**). In other words, white matter stimulation versus stimulation in the cortex had very different effects depending on if we stimulated in the cingulate, the lateral prefrontal lobe, or the lateral temporal lobe.

Dissecting this relationship further by relating the responses to the continuous distance to the grey-white boundary, we found three response categories: 1) overall increase with distance to the grey-white boundary (lateral prefrontal lobe), 2) overall decrease or no change (cingulate); 3) nonlinear u-shaped curve (lateral temporal lobe; **Fig. 5; Supplemental Figure 10, 11**). For instance, the relationship in the lateral temporal lobe between distance to the grey-white boundary and responses were better reflected by a u-shaped curve locally and distally with the best fit being a second order polynomial (average Pearson’s linear correlation: lateral temporal, local: 0.18±0.04, distant: 0.29±0.03; second order polynomial model fit significantly different to a constant term model; local: p<= 0.0102; distant: p<=0.0024; n=125 stimulation sites). In contrast, with stimulation in the lateral prefrontal lobe, the relationship was best described as a linear relationship across measures (average Pearson’s linear correlation: lateral prefrontal, local: 0.20±0.02, distant: 0.15±0.04). We found, in local sites, a linear relationship fit all four measures in the lateral prefrontal lobe best (F-test comparing model to a constant term model; p< 0.0005), but a second order polynomial best fit all the distant neural measures (F-test comparison to a constant term model; p< 0.00001; n=125 stimulation sites). In contrast, in the cingulate, we did not find a significant linear or nonlinear relationship between distance to the grey-white boundary and the neural responses, though there was an overall negative trend as we moved toward the white matter (average Pearson’s linear correlation: cingulate, local: - 0.12±0.09, distant: 0.08±0.05; F-test comparison fitted line to a constant term model; p>= 0.0461; n=105 stimulation sites; **Fig. 5; Supplemental Figure 11**). These complex relationships are apparent when the results are shown as a surface representation (**Fig. 5B**; **Supplemental Figure 12**).

**Figure 5.**
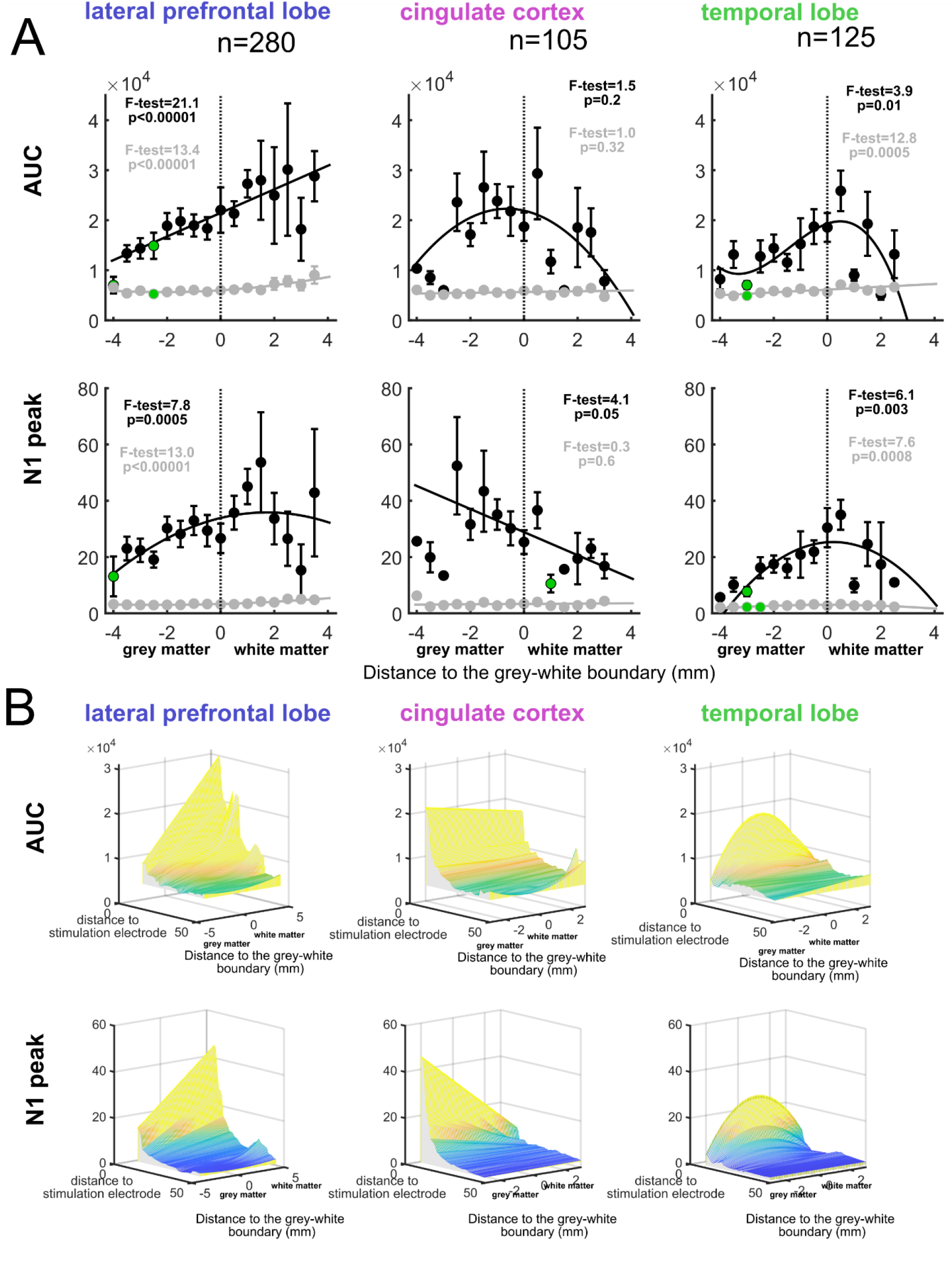
Region-specific effects on stimulation responses relative to grey and white matter neural responses in the lateral prefrontal lobe, lateral temporal lobe, and cingulate cortex. **A**. Local (<15mm, black) and distant (<15mm, grey) AUC and N1 peak responses plotted relative to the distance to the grey-white matter boundary for the different brain regions. Lines are GLM polynomial fits with the model order based on the minimum AIC value per fit. Each distance point and standard error bar has at least four stimulation sites contributing to each point. F-statistics and p-values for the model fits as compared to a model with a constant term for the local (black) and distant (grey) sites. **B**. Surface plots of GLM fitted lines for different stimulation to recording distances relative to the distance to the grey-white boundary for the N1 peak and AUC responses for the different brain regions.

### Orientation of the Stimulation Dipole Alters Responses

Not only could the location relative to grey and white matter be important, but we hypothesize that the orientation of the simulating dipole to neuronal structures could be crucial in sculpting a response [28,42]. We calculated the orientation of the bipolar pair of electrodes relative to the cortical column for both local and distant responses (**Fig. 6A; Supplemental Fig. 13**) with 0° indicating an alignment of the effective dipole with the axis of dendrites of pyramidal neurons in the grey matter. We found a rise in the local peak, AUC, N1 peak, and N2 peak responses with increasing angles (orientation of the electrode relative to the nearest cortical axis) up until 90° and falling off again as the orientation moved toward 180 ° (**Fig. 6B; Supplemental Fig. 13**). This local response relationship was best modeled with a third-degree polynomial fit. In contrast, the angle of orientation had no impact on distant responses. Neither the local or distant orientation vs. responses were linear (mean Pearson’s linear correlation rho for all voltage measures=0.05±0.08; N=52).

**Figure 6.**
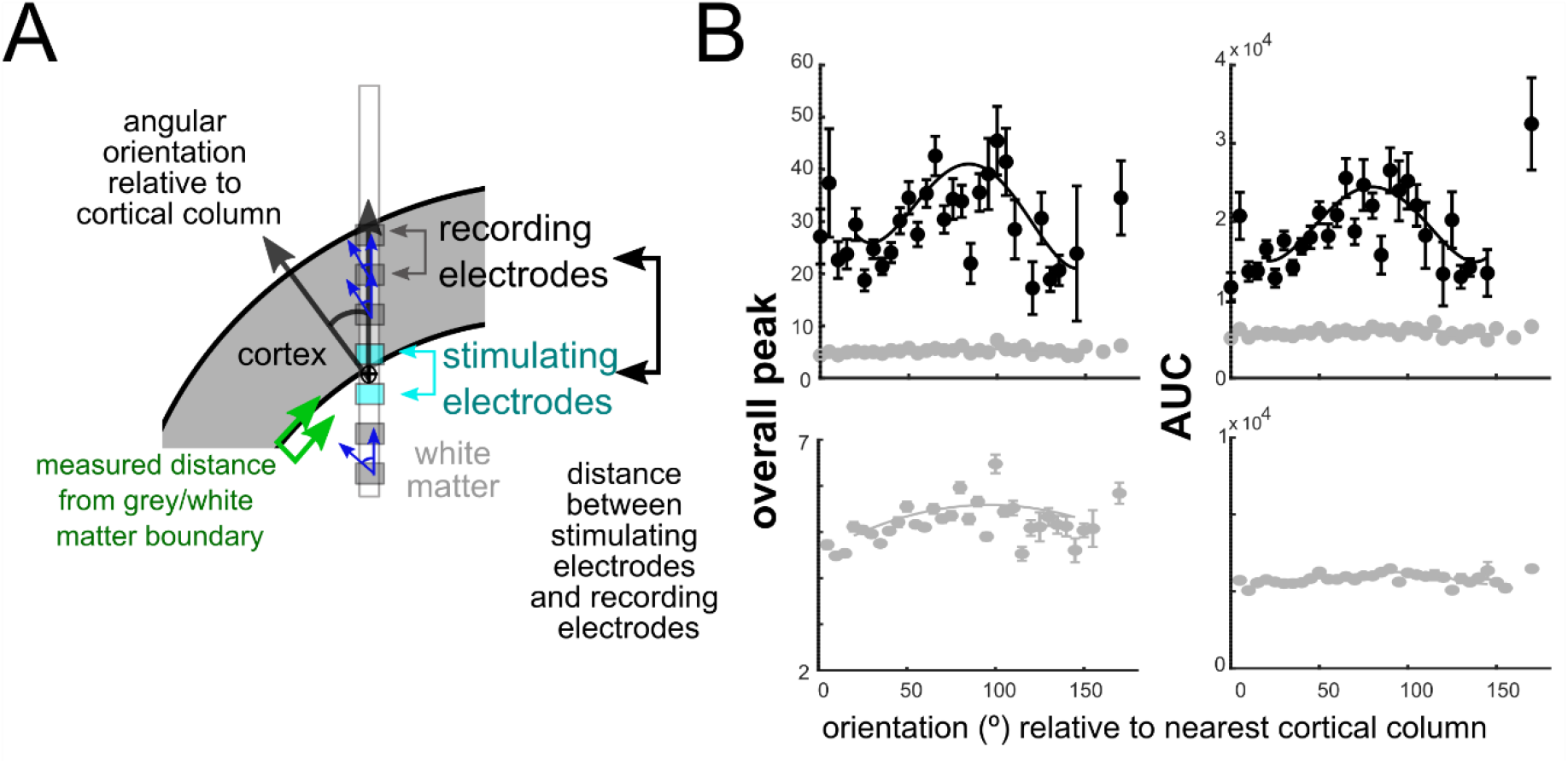
Relationship between electrode orientation and response. **A**. Measures of orientation (°) of the bipolar pair of electrodes to the cortical axis. **B**. Local (<15 mm, top row) and distant (>15 mm, top and bottom row) overall peak and AUC responses plotted relative to the orientation (°) of the bipolar pair of electrodes to the cortical axis. Individual points and error bars are mean and standard error. Data is from all contacts, not just the above-threshold responses.

As orientation could also be dependent on whether the stimulating bipolar pair of electrodes was in grey matter versus white matter, we subdivided the data based upon whether the stimulating electrode was in the white matter or cortical volume (**Supplemental Fig. 13**). We found the relationship between bipolar pair orientation versus responses to be linear for local recording sites, reaching significance for the overall peak and AUC (peak, local: rho=0.56; p=0.0048; AUC: local: rho=0.61; p=0.002; N1 peak, local: rho=0.46; p=0.03; N2 peak: local: rho=0.46; p=0.02; N=52). Yet, the correlation between cortical orientation and responses were small or nonexistent for the local recording sites when the electrode pair was in the white matter (peak, local: rho=0.09; p=0.63; AUC: local: rho=0.06; p=0.78; N1 peak, local: rho=0.17; p=0.39; N2 peak: local: rho=0.22; p=0.26; N=52). This result makes sense considering the stimulation sites in the white matter far away from the cortical dipole will have less of a correlation with stimulation orientation relative to the cortex. Interestingly, in contrast, responses at distant recording sites (>15 mm) had a had no clear relationship with orientation (peak, distant: rho=-0.23; p=0.22; AUC: distant: rho=-0.13; p=0.49; N1 peak, distant: rho=-0.21; p=0.24; N2 peak: distant: rho=-0.22; p=0.22; N=52; **Supplemental Fig. 13**). In other words, for stimulation in white matter and distant contacts, orientation may not play as much of a role as it does for local cortical stimulation.

## Discussion

Through a systematic DES study in a large intracranial data set we found evidence for basic rules of intracranial stimulation in the human brain (**Fig. 7**). We found that, as expected, increasing stimulation duration leads to larger responses at any given location. In addition, unlike previous studies involving trains of stimulation [24], we found that single pulse responses vary non-linearly with injected current and this is true both nearby the stimulation and at a distance. Specifically, increasing stimulation from 1 to ∼5 mA linearly increases the response but that beyond ∼5 mA the response either plateaus or even decreases. We also confirmed that decreasing stimulation distance to white matter can increase distant responses while the largest local responses occurred with stimulation at the grey-white boundary across brain regions while distant responses were best induced with stimulation entirely in white matter (**Fig. 7**).

**Figure 7.**
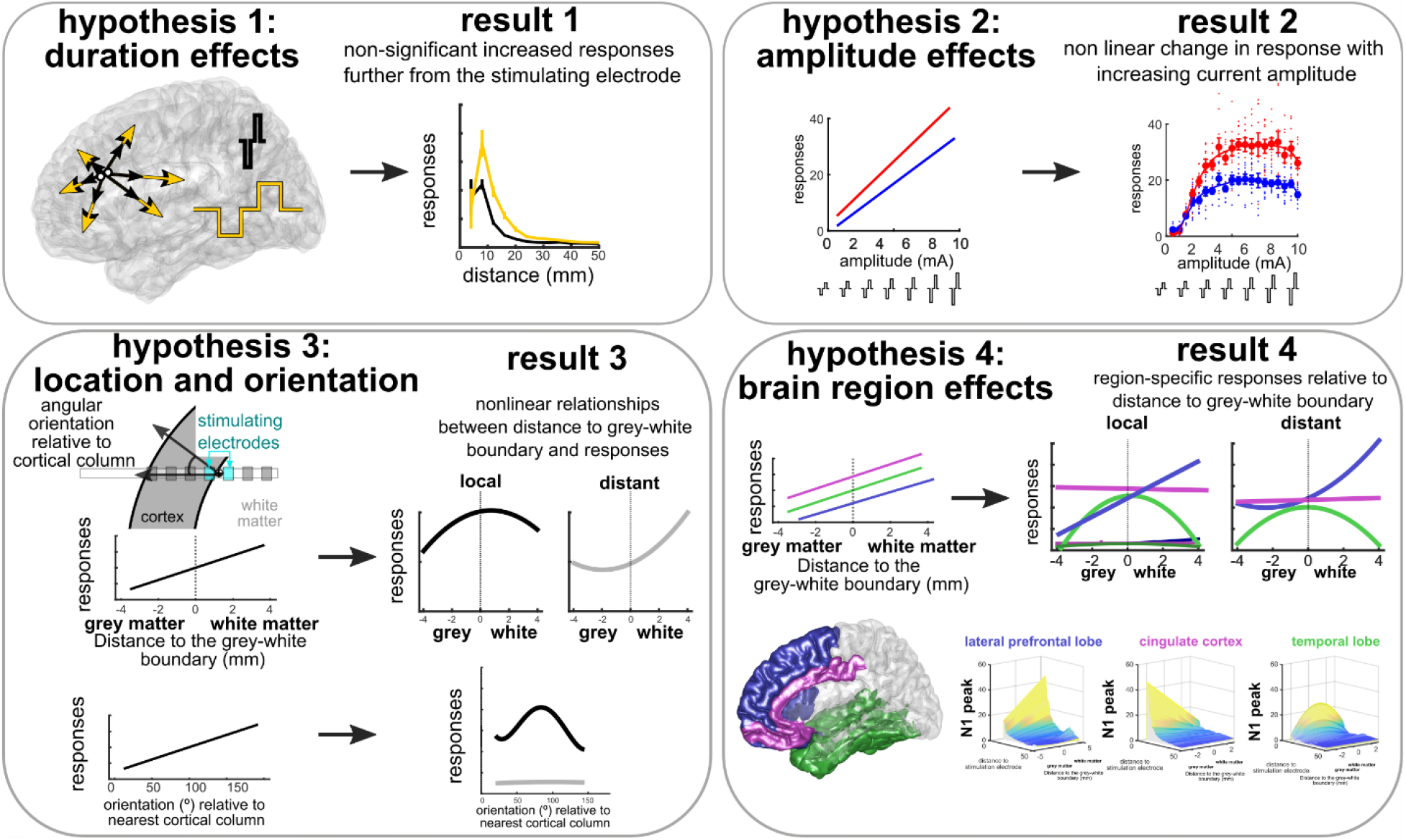
Summary of hypotheses and relevant results. Summary of hypotheses and relevant results.

We also found, as hypothesized, that increasing duration increased the neural responses both locally and distally, though, as we had only two duration steps, we could not determine if the relationship was linear. Related companion work found there were no differences in responses for intracranial sEEG responses for two larger duration steps (0.5 ms and 1 ms) which could indicate there is a nonlinear saturation effect with changing duration, though, interestingly, there were differences in the response on the scalp as recorded by high density EEG shown by Parmigiani et al. (a companion manuscript in preparation/bioRxiV). Future work may not only need to include more duration steps, but also to identify if duration and amplitude are independent variables or merely parameters controlling total current which is what really dictates response, particularly if there are regional differences with changing waveform duration.

Further, the stimulation parameter effect depended on what aspect of the voltage waveform was measured. For input current, for example, N1 peak and N2 peak exhibited different saturation points. This difference likely relates to differences in N1 and N2 mechanisms; the N1 peak is thought to reflect local excitation while N2 relates to a network polysynaptic response or inhibitory rebound (N2, [65,88,89]). Indeed, the decrease in the response at the highest amplitudes could imply that there is a neural suppression through saturation or the activation of inhibitory circuits in nearby cortical regions. In terms of therapy and clinical mapping, then, the question whether there is a reason for going to high amplitudes beyond 7 mA unless there is another physiological response desired such as inducing seizures or completely suppressing local neural activity [90], particularly with safety considerations [91].

In contrast to our results here, in prior work, we demonstrated a consistent linear relationship between responses and stimulus current from trains of stimulation [24], highlighting an important distinction between SPES and train-based stimulation. It is possible that repeated pulses act to engage larger reverberating or oscillating networks with increasing amplitude while the mechanisms underlying responses to SPES can reach a limit by not engaging these widespread networks. Further tests, including the use of microelectrode recordings to parse individual neuronal activity during these different stimulation approaches [92].

With this large data set and consistent, systematic approach, we found that stimulation at the grey-white boundary and especially white matter resulting in the largest responses (**Fig. 7**). This effect appears most striking in the lateral prefrontal cortex. Strikingly, similar results were demonstrated by a companion study conducted by a companion study by Parmigiani et al., particularly with a stronger effect of white matter stimulation increasing the N2 peak intracranially and both the N1 and N2 peak as recorded on the scalp by high density EEG (in preparation/bioRxiV). We also found a stronger effect on responses with the bipolar electrode oriented perpendicular to the closest cortical axis which is in keeping with results from high density EEG data collected by Parmigiani et al. (in preparation/bioRxiV). The results presented here are interestingly in light of recent work where trains of stimulation were more effective at decreasing high frequency activity when stimulation was in grey matter as opposed to white matter, with stronger effects in the neocortex versus the medial temporal lobe [28]. Further work is needed to resolve the differences between low frequency voltage responses such as CCEP, high frequency responses, and the types of stimulation. Regardless, the fact that different regions show different location dependency likely reflects the different regional microarchitecture (e.g. arrangement and number of particular neuronal types) as well as differential network connectivity (e.g. the number of long-range connections from the region).

It is possible that white matter stimulation is more effective because direct axonal stimulation can result in both antidromic and orthodromic conduction [93,94]. Congruent with this finding is the targeting of white matter tracts and axonal stimulation to treat depression and Parkinson’s, [58,62]. Another key possibility is that there could be network effects engaging thalamic networks [82] which could explain why stimulation in white matter increased N2 peak and AUC measures (which are generally the slower portions of the responses) but had less of an effect on overall peak and N1 peak responses (which are faster responses and could be more locally generated). Interestingly, in our study the local responses had a peak response with stimulation at the grey-white boundary, not necessarily in white matter itself, particularly in the lateral temporal lobe. We posit there could be an effect of being closer to the axon hillocks of large pyramidal cells in cortical layers 4-6 and that the grey-white boundary is a convergence point for multiple output neurons or is a site with a higher concentration of excitatory versus inhibitory contributions which could explain the peak local responses [83]. To test these ideas, modelling in combination with cytoarchitectonic maps are likely necessary [84,85] in addition to sampling of neural data on microscale levels [14,92,95].

For all these comparisons, we focused on distances between sites based on the geometry of the brain structures. This choice was made to limit the scope to a purely distance measure. While there are exciting conclusions to be made relative to white matter tractography, connectivity, and stimulation [29,60], this data set did not have consistent enough sampling of noninvasive scans mapping white matter (such as diffusion tensor imaging, or DTI). Further, our approach and a companion study by Parmigiani et al. (in preparation, bioRxiV) was to start from a naïve perspective such that this hypothesis testing and results using geometry could generate independent conclusions which, in future studies, can be mapped to structural and functional connectivity [29,60]. For example, Human Brain Connectome data could be instrumental in answering this question [87]. Alternatively, WM tractography data might be more available in the future with the possible increase in clinical usefulness of white matter tractography as a clinical tool, such as to help understand epilepsy networks [96,97].

These conclusions must be tempered by an awareness that our sampling of the brain remains sparse in two respects. First, at maximum we are recording from ∼200 locations with most sites in frontal and lateral temporal areas, many in white matter and often separated by larger distances. Nuances of the effects of stimulation at different distances and at different subregions of the brain might be lost. Second, this also means our stimulation sites are not uniformly distributed across the brain. Because of clinical constraints, we have few stimulation examples from primary sensory or motor cortex or most subcortical structures and those regions may react different to stimulation or respond differently to stimulation elsewhere. Moreover, bipolar stimulation through macroelectrode contacts means that the electrical field generated is relatively large, spanning millimeters [6]. This large size of the field makes our localization along the grey-white continuum imprecise which could be complicated by possible brain shift [69,98] though we focused on sEEG electrodes which are less prone to brain shift compared to subdural grids and strips [99]. None-the-less, even with these caveats we would expect that the relationship we are seeing with respect to grey-white location to hold. Another important caveat is that there is likely to be an effect of brain state on the stimulation-response input-output relationship [60,100]. While we focused on data gathered while the participants were awake it is possible that during sleep, for example, the stimulation rules may be different [101].

Finally, of course, there are always questions surrounding conclusions drawn from findings obtained in the setting of pathology – in this case epilepsy. Epilepsy is clearly a network process and this could have an impact on the stimulation input/output relationship [102,103]. Medications could also alter the physiology. We attempted to avoid these potential confounds by excluding channels from either recording or stimulation which showed substantial epileptiform activity. In addition, by looking across many locations in many patients on different medications and with different etiologies, we should be ‘averaging out’ the influence of pathological since the pathology will be unique to individual patients.

This work, alongside a growing number of systematic studies of DES and DBS [27,28,51,58,104–107] suggests that it may be possible to identify consistent stimulation parameter – output rules enabling a determination of brain region, where in the grey-white matter and with what amplitude, duration, and frequency a stimulus should be given to produce a specific local and distant output. This would be of immense assistance in developing targeted, effective, treatments for a wide range of neuropsychological challenges [14,24,60,108]. Therefore, the main results of grey and white matter stimulation and local and distant effects in our study suggest that therapies using stimulation should be modified per targeted neurophysiological outcome. For instance, it might be worthwhile to target stimulation across the grey-white junction when large, local responses are needed to reach a neurophysiological and behaviorally relevant therapeutic goal. Alternately, if a therapeutic goal is to induce widespread, network-level changes which may be ideal in treating certain forms of epilepsy [109] or mapping network circuits [29,49,52], then it may be preferable to target white matter in the lateral prefrontal cortex. Conversely, targeting grey matter such as in the cingulate might be useful when highly localized responses or inducing small circuit changes are needed such as in using microstimulation to impact memory formation [14,110]. Indeed, instead of proposing a “one stimulation parameter set fits all” approach, tailored therapeutic treatments driven by a growing body of knowledge of the complex, but knowable and mappable, relationship of stimulation parameters and different types of targeted neural dynamics may provide the most reliable DES treatments for individual patients.

## Abbreviations

SPES: Single pulse electrical stimulation
CCEP: cortico-cortico evoked potential
DES: Direct electrical stimulation
EEG: electroencephalogram
sEEG: stereo EEG
ERP: event-related potential
LFP: local field potential
ELA: electrode labeling algorithm
AUC: area under the curve
OCD: obsessive compulsive disorder

## Acknowledgments

We would like to thank Giovanni Piantoni, Jean-Baptiste Eichenlaub, Erica Johnson, Gavin Belok, Mia Borzello, Kara Farnes, Gavin Belok, Dan Soper, Constantin Krempp, Jaquelin Dezha-Peralta, and Pariya Salami for help in data collection. We would like to especially thank the patients for participating in the study.

