## Supplemental Data for "Impact of stimulation location relative to grey and white matter on single pulse electrical stimulation responses in the human brain"

Formatted: Line spacing: single

**Supplemental data**

**Supplemental Figure 1. Counts of stimulation sites relative to brain regions and stimulation tests across the data set.**

**Supplemental Figure 2. Example stimulation location relative to stimulation spread.**

**Supplemental Figure 3. Interaction between responses and distances between stimulation sites and recording sites.**

**Supplemental Figure 4. Interaction between responses and distances between stimulation sites and recording sites taking into account contacts along the same depth as the stimulating electrode and not along the same depth.**

**Supplemental Figure 5. Single pulse stimulation response depends on location relative to cortex, white matter, and subcortical regions.**

**Supplemental Figure 6. Region-specific effects on stimulation responses relative to grey and white matter neural responses in the lateral occipital lobe, the parietal lobe, and subcortical regions.**

**Supplemental Figure 7. Distances and orientations of stimulating electrodes relative to major landmarks.**

**Supplemental Figure 8. Mapped surfaces of the fitted responses using GLM and linear (first order) and quadratic (second order) models.**

**Supplemental Figure 9. Single pulse stimulation response depends on location relative to cortex and white matter as well as brain region.**

**Supplemental Figure 10. Single pulse stimulation response depends on location relative to cortex and white matter as well as brain region.**

**Supplemental Figure 11. Participant (each color is a different participant) average responses with stimulation sites relative to the distance to the grey-white boundary.**

**Supplemental Figure 12. Mapped surfaces of the fitted responses using GLM fits for the lateral prefrontal lobe, the cingulate, the lateral temporal lobe, and subcortical structures.**

**Supplemental Figure 13. Distances and orientations of stimulating electrodes relative to major landmarks.**

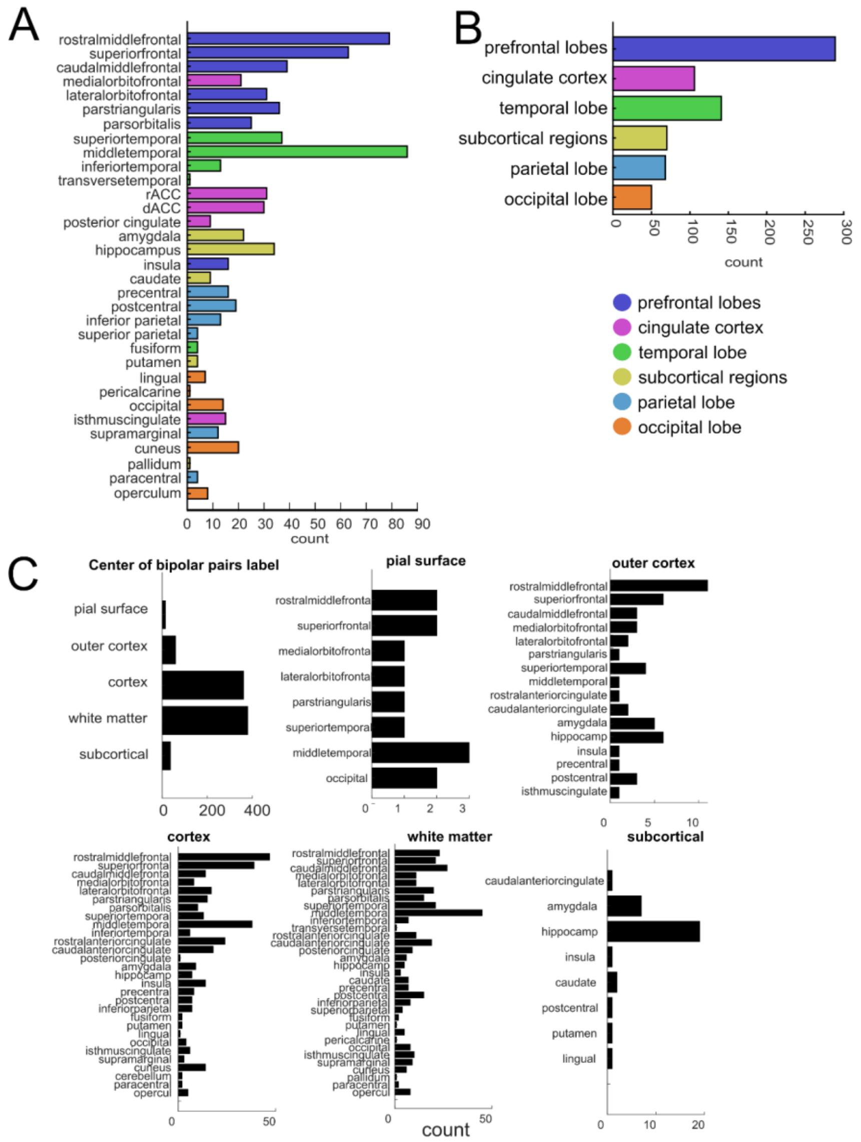


**Supplemental Figure 1. Counts of stimulation sites relative to brain regions and stimulation tests across the data set.** Distribution of center points of electrodes and the nearby brain regions in a bipolar pair outside the pial surface, outer cortex, cortex, white matter, or located in subcortical structures, N=52.

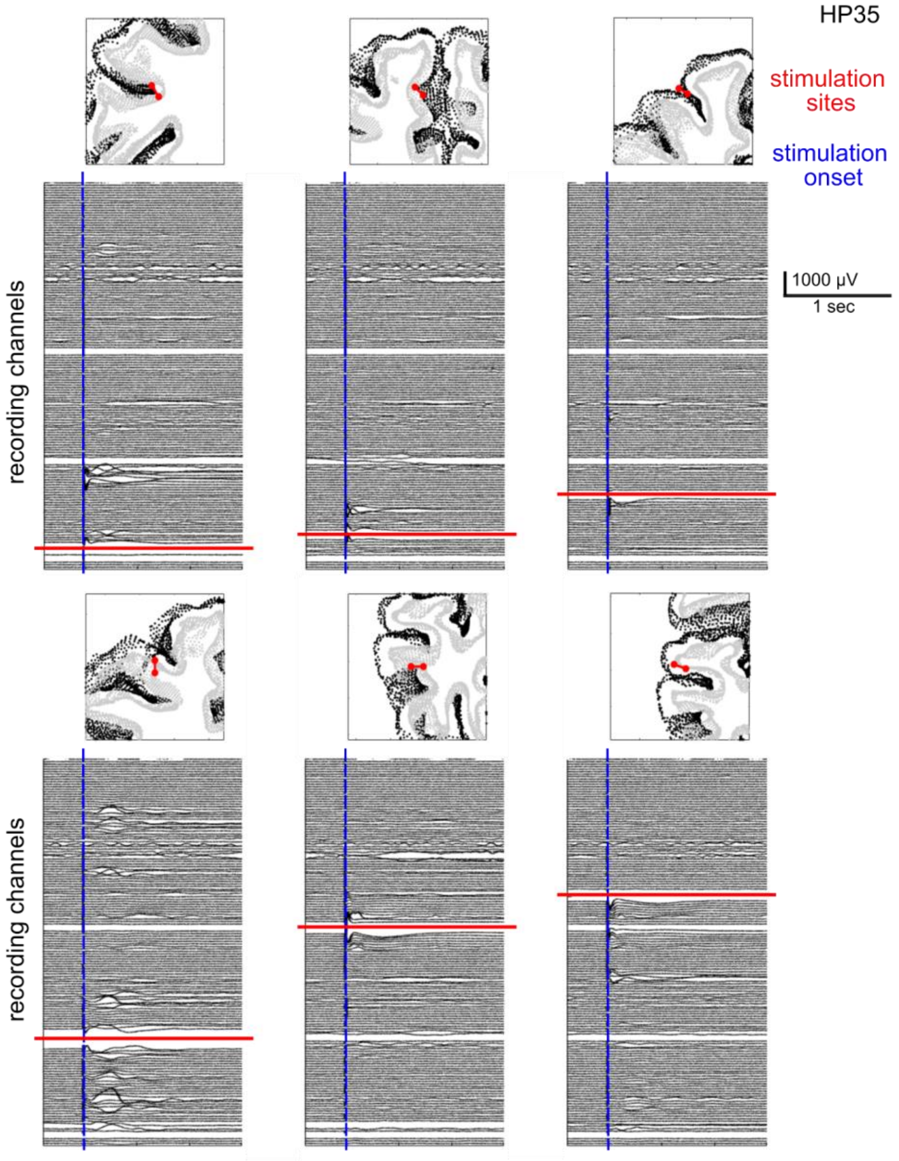

**Supplemental Figure 2. Example stimulation location relative to stimulation spread. Participant: HP35.** Channel order shown in order of original recording following bipolar re-referencing. Red lines indicate the relative location of the stimulating electrodes to the recording electrodes (in black lines). Blue lines indicate stimulation onset. In the panels above the traces are images of the locations of the stimulating electrodes in coronal slices of the participant, with black dots indicating points of the pial surface and grey dots indicating the grey matter-white matter boundary.

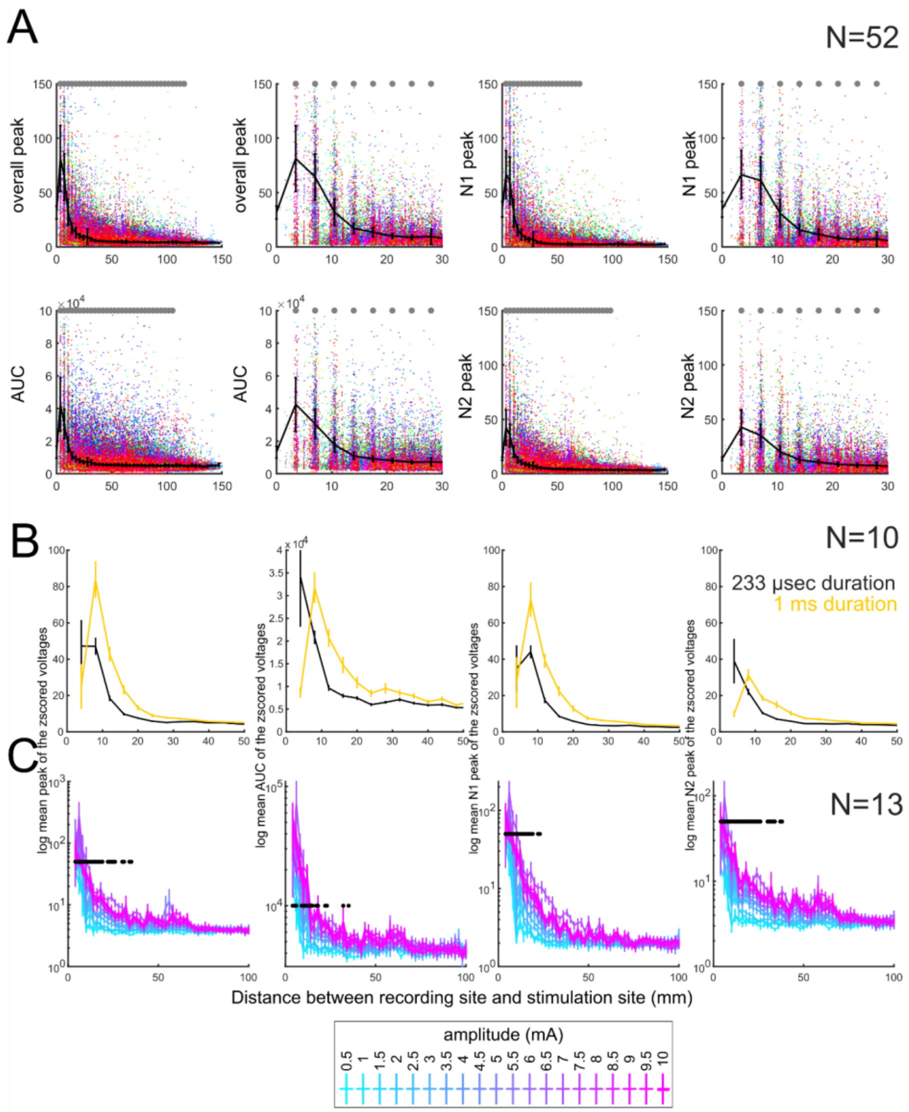


**Supplemental Figure 3. Interaction between responses and distances between stimulation sites and recording sites. A.** Distance versus responses for the overall peak, AUC, N1 peak, and N2 peak responses, dots with color coding representing different participants. Top row: overall peak and N1 peak, with the left plots per measure the entire distance range and the right plots per measure 0 and 30 mm scales. Bottom row: AUC and N2 peak, with the left plots per measure the entire distance range and the right plots per measure 0 and 30 mm scales. The black error bars are mean and standard deviation of the distributions across participants. **B**. Effect of increasing duration from 233 µsec (black) to 1 ms (green) on spread of the neural response in N=9 participants in the overall peak, AUC, N1 peak, and N2 peak responses. The difference is not significant (not below p<0.0063, corrected for multiple comparisons; Wilcoxon rank-sum test) between 233 µsec (black) and 1 ms (green) stimulation durations at each distance step. **C**. Effect of increasing amplitude from 0.5 (cyan) to 10 mA (magenta) on spread of the neural response in N=13 participants in the overall peak, AUC, N1 peak, and N2 peak responses. The difference is significant (p<0.00005; Kruskal-Wallis multiple comparisons test) between the different amplitude steps at stimulation-recording electrode distance steps <30 mm as indicated by the black dots overlaid on the traces.

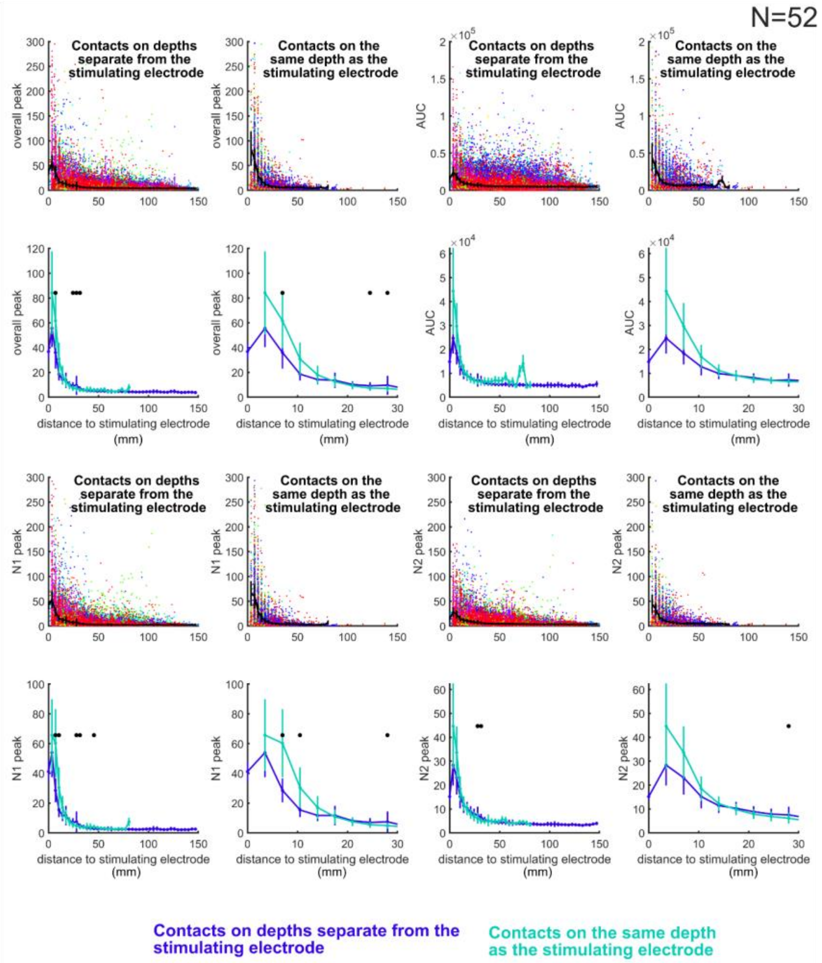


z

**Supplemental Figure 4. Interaction between responses and distances between stimulation sites and recording sites taking into account contacts along the same depth as the stimulating electrode and not along the same depth.** Distance versus responses for the overall peak (upper left four plots), AUC (upper right four plots), N1 peak (lower left four plots), and N2 peak responses (lower right four plots) In each quadrant, the two top rows have dots with color coding representing different participants, with plots of the responses with distance depending on if the recording electrode was on the same depth shaft (right) or not (left). The black error bars are mean and standard deviation of the distributions across participants. Bottom plots of error bars are the mean and standard deviation of the distributions across participants for the recording electrodes along the same shaft (cyan) or different electrode shafts (royal blue), with the zoomed out view (left) and 0 and 30 mm range (right). The difference is significant ($p<0.00005$; Wilcoxon rank sum test) between the contacts along the same depth as the stimulating electrode versus not as indicated by the black dots overlaid on the traces.

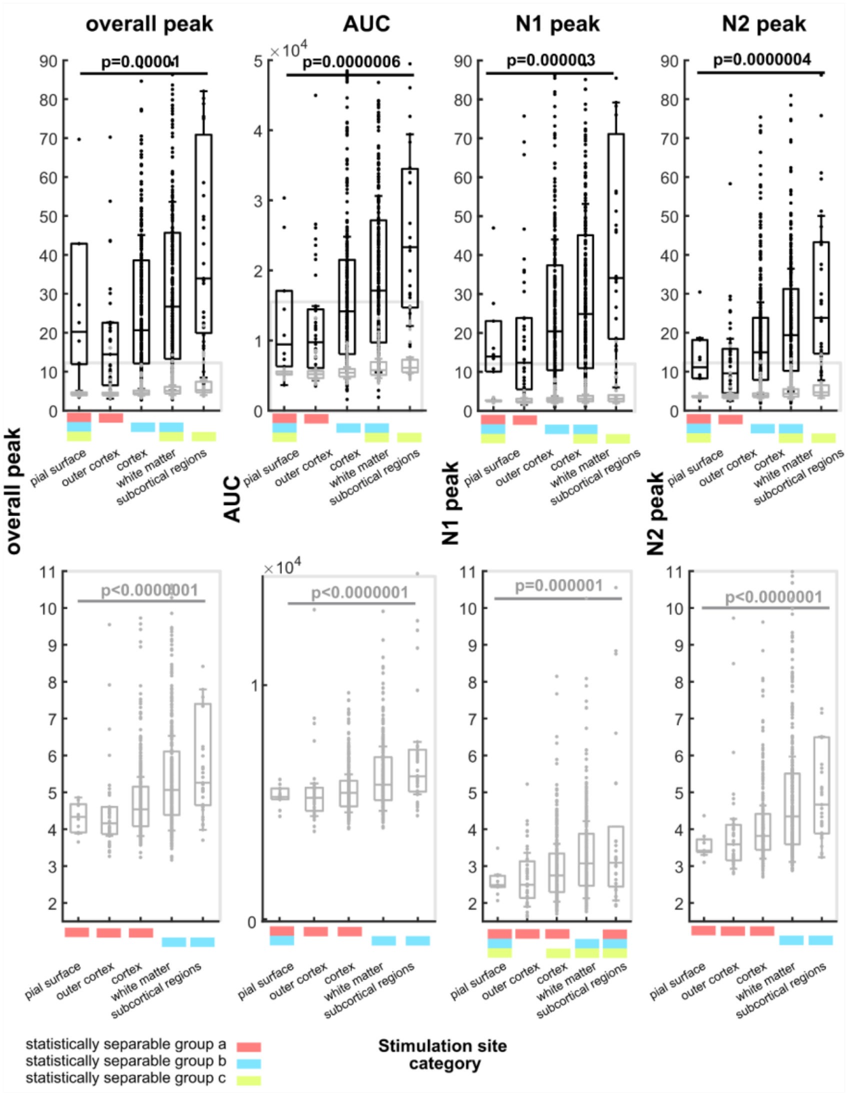


**Supplemental Figure 5. Single pulse stimulation response depends on location relative to cortex, white matter, and subcortical regions.** Effect of stimulation site on peak neural responses (1st panel), area under the curve (AUC; 2nd panel); N1 peak (3rd panel) or N2 peak (4th panel). Recording sites are separated into local (<15mm away from the stimulation site;

white bars with black outlines, top row of figures) or distant sites (recording sites >15mm away from the stimulation site, grey outlined bars, and expanded in the bottom row). N=52; p-values are from Kruskal-Wallis multiple comparisons tests, followed by a *post hoc* Tukey-Kramer test of significance. Color bars below each category is to indicate statistically separable groups resulting from a pairwise comparison of each group to every other group with the *post hoc* Tukey-Kramer test of significance. In this case, the regional categories (e.g. cortex or white matter) shown with red bars are not significantly different from one another but are significantly different to the regional categories with only blue or green bars.

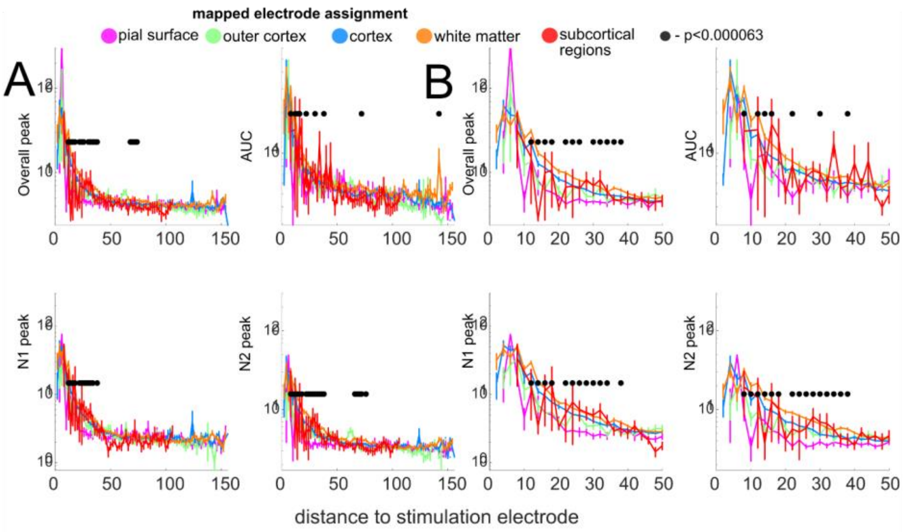


**Supplemental Figure 6. Region-specific effects on stimulation responses relative to grey and white matter neural responses in the lateral occipital lobe, the parietal lobe, and subcortical regions. A**. Effect of changing stimulation location relative to the cortex and white matter volumes on responses and distance relative to the stimulation site at 7mA and 233 µsec in N=52 participants in the overall peak, AUC, N1 peak, and N2 peak responses. The difference is significant (p<0.00005; Kruskal-Wallis test) between the different locations at stimulation-recording electrode distance steps <50 mm as indicated by the black dots overlaid on the traces. **B**. Same data but focusing on the local responses.

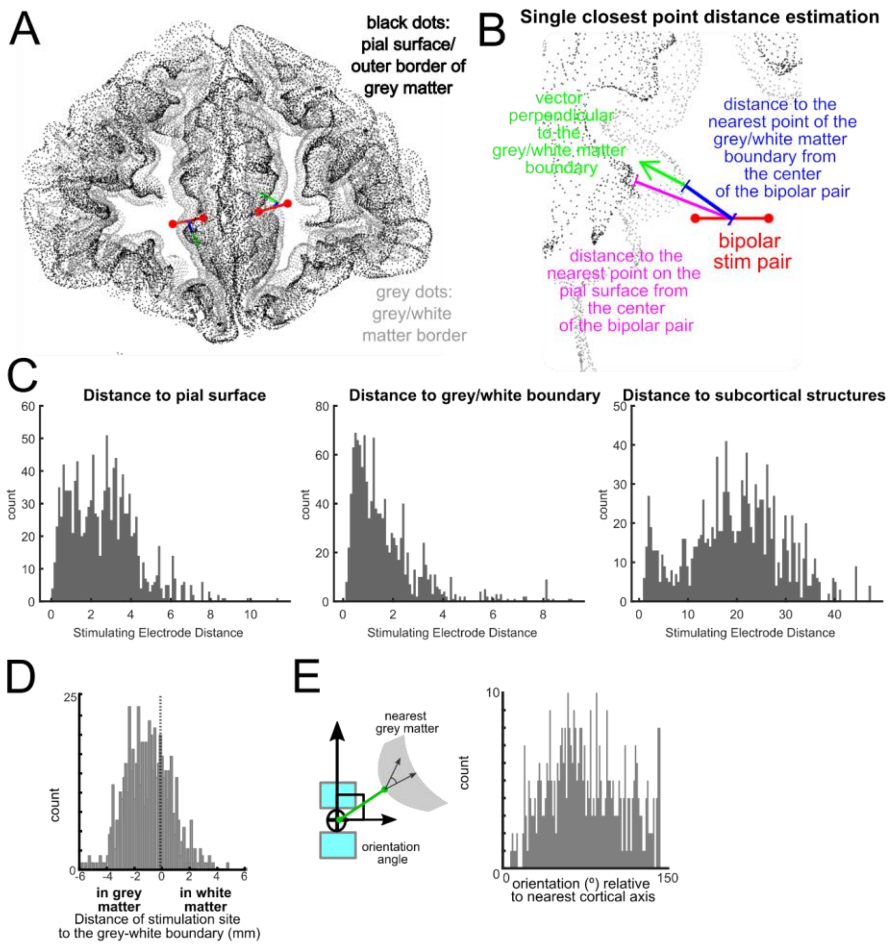


**Supplemental Figure 7. Distances and orientations of stimulating electrodes relative to major landmarks. A-B**. Location of two example bipolar pairs of electrodes relative to the grey and white matter in a single scanned slice, with an enlarged view of the different measurements which include the closest grey matter and white matter points from the center of the stimulating bipolar pair. **C**. Distribution of distances to the pial surface, the grey/white boundary, and to subcortical structures for all stimulation sites. **D**. Distribution of distances relative to the grey/white boundary across the data set. **E**. Orientation measure relative to the orientation of the closest cortical column and distribution of angles (º) across the data set and the distribution of orientations across all stimulation sites.

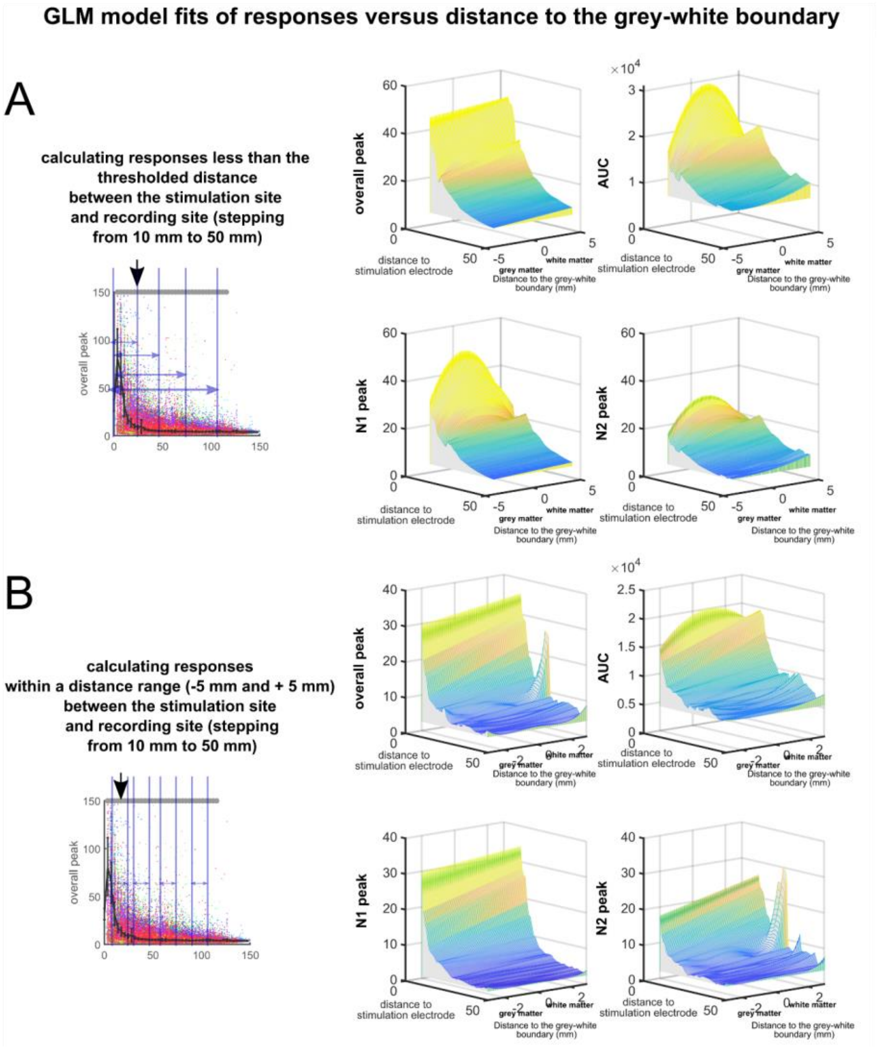


**Supplemental Figure 8. Mapped surfaces of the fitted responses using GLM and linear (first order) and quadratic (second order) models. A.** The GLM fit between each of the responses (overall peak, AUC, N1 peak, and N2 peak) and the distance to the grey-white boundary as we include electrodes further and further away from the stimulating electrode plotted as a surface. At each included electrode distance, the best GLM fit was identified (by minimizing the AIC value) to determine if the fit was best mapped to a linear model, a quadratic second order model, or a third order polynomial fit. **B.** The GLM fit between responses (overall peak, AUC, N1 peak, and N2 peak) and the distance to the grey-white boundary at each stepwise binned distance from the stimulating electrode (-5 to +5 mm) from the stimulating electrode plotted as a surface. At each included electrode distance, the best GLM fit was identified (by minimizing the AIC value) to determine if the fit was best mapped to a linear model, a quadratic second order model, or a third order polynomial fit..

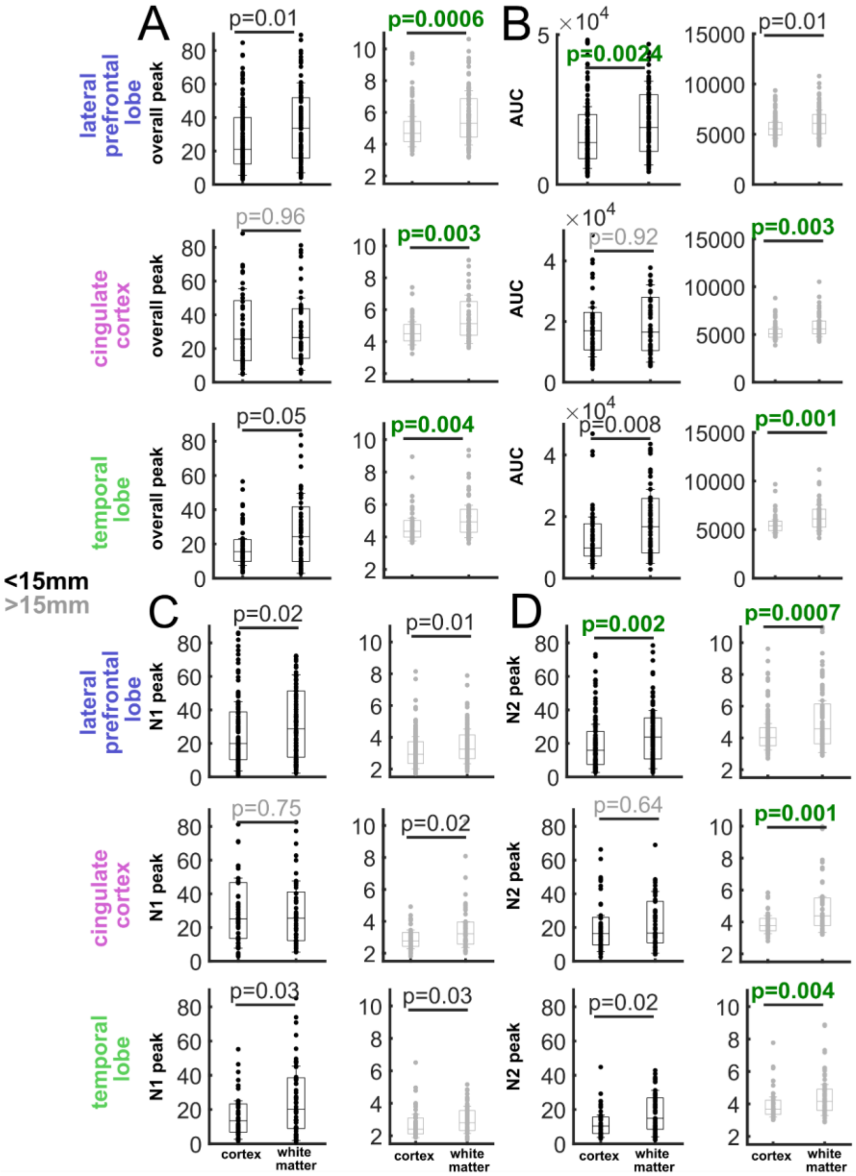


**Supplemental Figure 9. Single pulse stimulation response depends on location relative to cortex and white matter as well as brain region.** Effect of stimulation site on overall peak neural responses (A), area under the curve (AUC; B); N1 peak (C) or N2 peak (D) for the lateral prefrontal lobe (top row), cingulate cortex (middle row) and temporal lobe (bottom row). Recording sites are separated into local (<15mm away from the stimulation site; white bars with black outlines, left columns) or distant sites (recording sites >15mm away from the stimulation site, grey outlined bars, right columns). N=52; p-values are from Wilcoxon rank-sum tests comparing white matter versus cortical stimulation.

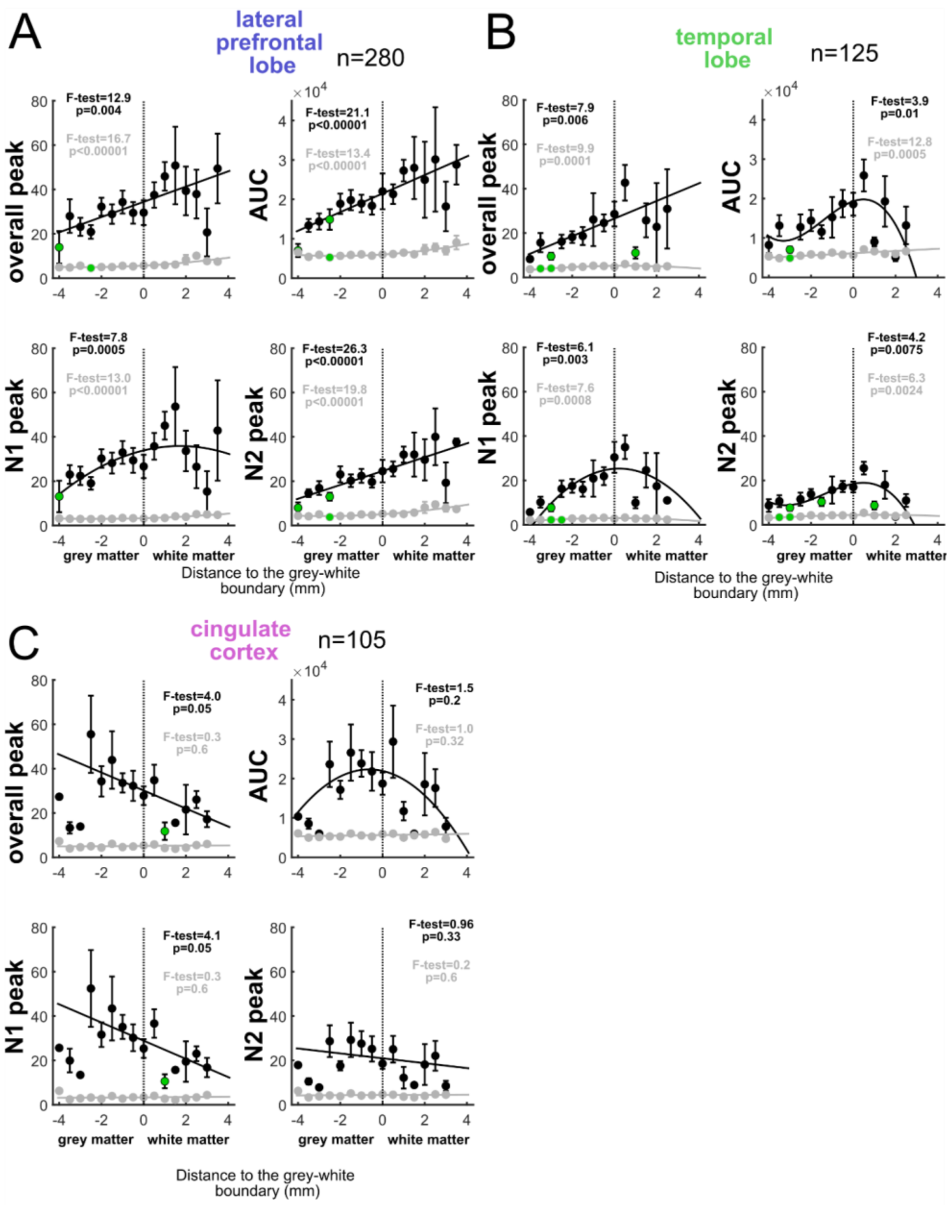

A
lateral prefrontal lobe
n=280
B
temporal lobe
n=125
C
cingulate cortex
n=105
overall peak
AUC
N1 peak
N2 peak
F-test=12.9 p=0.004
F-test=16.7 p<0.00001
F-test=21.1 p<0.00001
F-test=13.4 p<0.00001
F-test=7.8 p=0.0005
F-test=13.0 p<0.00001
F-test=26.3 p<0.00001
F-test=19.8 p<0.00001
F-test=7.9 p=0.006
F-test=9.9 p=0.0001
F-test=3.9 p=0.01
F-test=12.8 p=0.0005
F-test=6.1 p=0.003
F-test=7.6 p=0.0008
F-test=4.2 p=0.0075
F-test=6.3 p=0.0024
F-test=4.0 p=0.05
F-test=0.3 p=0.6
F-test=1.5 p=0.2
F-test=1.0 p=0.32
F-test=4.1 p=0.05
F-test=0.3 p=0.6
F-test=0.96 p=0.33
F-test=0.2 p=0.6
grey matter
white matter
Distance to the grey-white boundary (mm)

**Supplemental Figure 10. Single pulse stimulation response depends on location relative to cortex and white matter as well as brain region. A-C**. Local (<15 mm, black lines and curves, left) and distant (>15 mm, grey lines and curves, expanded to the right) peak in the 1 second after stimulation (top left), AUC (bottom left), N1 peak (top right), and N2 peak (bottom right) responses plotted relative to the distance to the grey-white matter boundary for the lateral prefrontal lobe (**A**), lateral temporal lobe (**B**), and cingulate (**C**). Data is from all contacts per brain region, not just the above-threshold responses. Green dots- responses at different distances to the grey-white boundary significantly different to responses at 0 mm same response measure, $p<0.000062$ (Wilcoxon rank sum test; multiple comparisons corrected). F-statistics and p-values for the model fits as compared to a model with a constant term for the local (black print) and distant (grey print) sites.

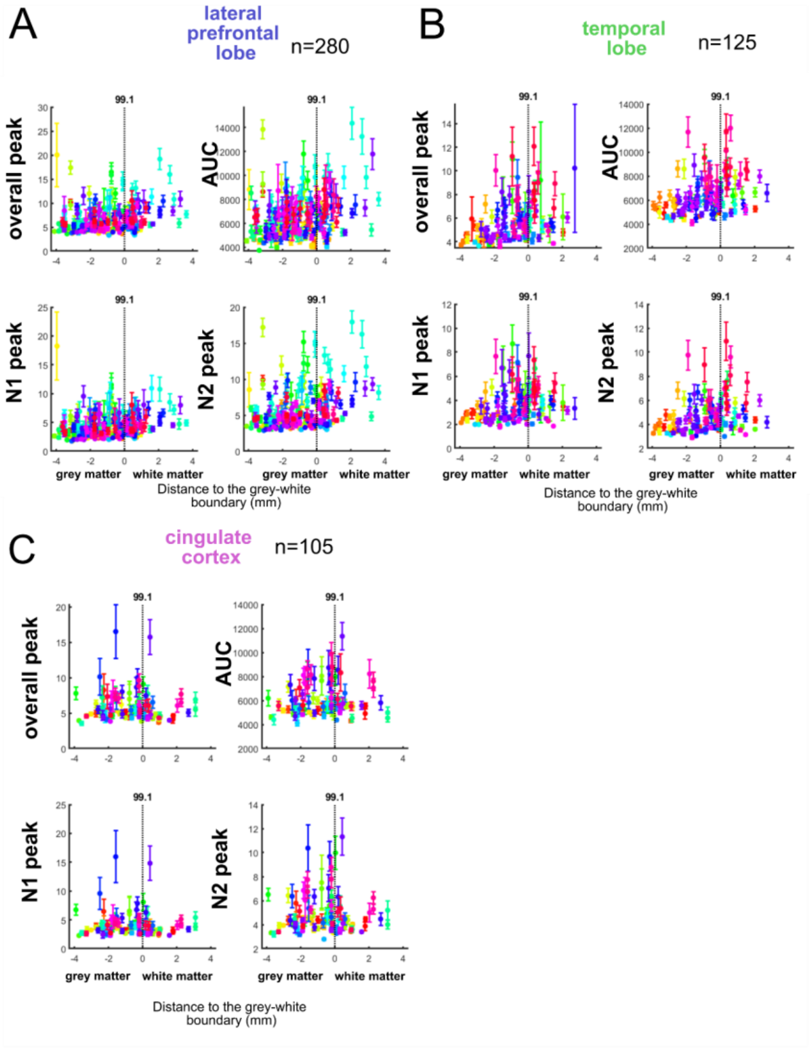


**Supplemental Figure 11. Participant (each color is a different participant) average responses with stimulation sites relative to the distance to the grey-white boundary.** Distant (<15mm) peak, AUC, N1 peak, and N2 peak responses plotted relative to the distance to the grey-white matter boundary for the different brain regions per participant. Each distance point and standard error bar is a stimulation point with n>=10 trials contributing to each point.

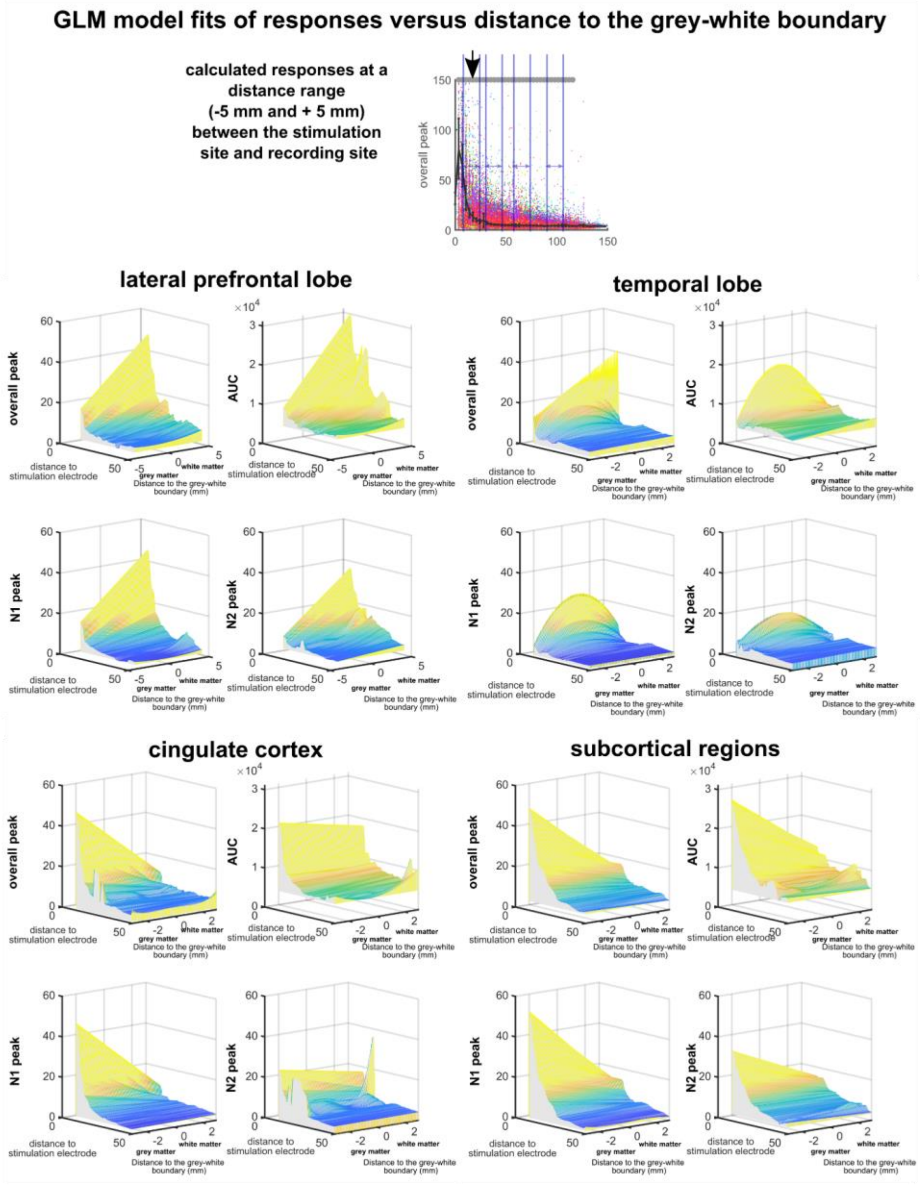
GLM model fits of responses versus distance to the grey-white boundary
calculated responses at a distance range (-5 mm and + 5 mm) between the stimulation site and recording site
overall peak
lateral prefrontal lobe
temporal lobe
cingulate cortex
subcortical regions
AUC
N1 peak
N2 peak
distance to stimulation electrode
grey matter
white matter
Distance to the grey-white boundary (mm)

**Supplemental Figure 12. Mapped surfaces of the fitted responses using GLM fits for the lateral prefrontal lobe, the cingulate, the lateral temporal lobe, and subcortical structures.** The GLM fit between each of the responses (overall peak, AUC, N1 peak, and N2 peak) and the distance to the grey-white boundary as we include electrodes further and further away from the stimulating electrode plotted as a surface. At each included electrode distance, the best GLM fit was identified (by the minimum AIC value per model order) to determine if the fit was best mapped to a linear model, a quadratic second order model, or a third order polynomial model.

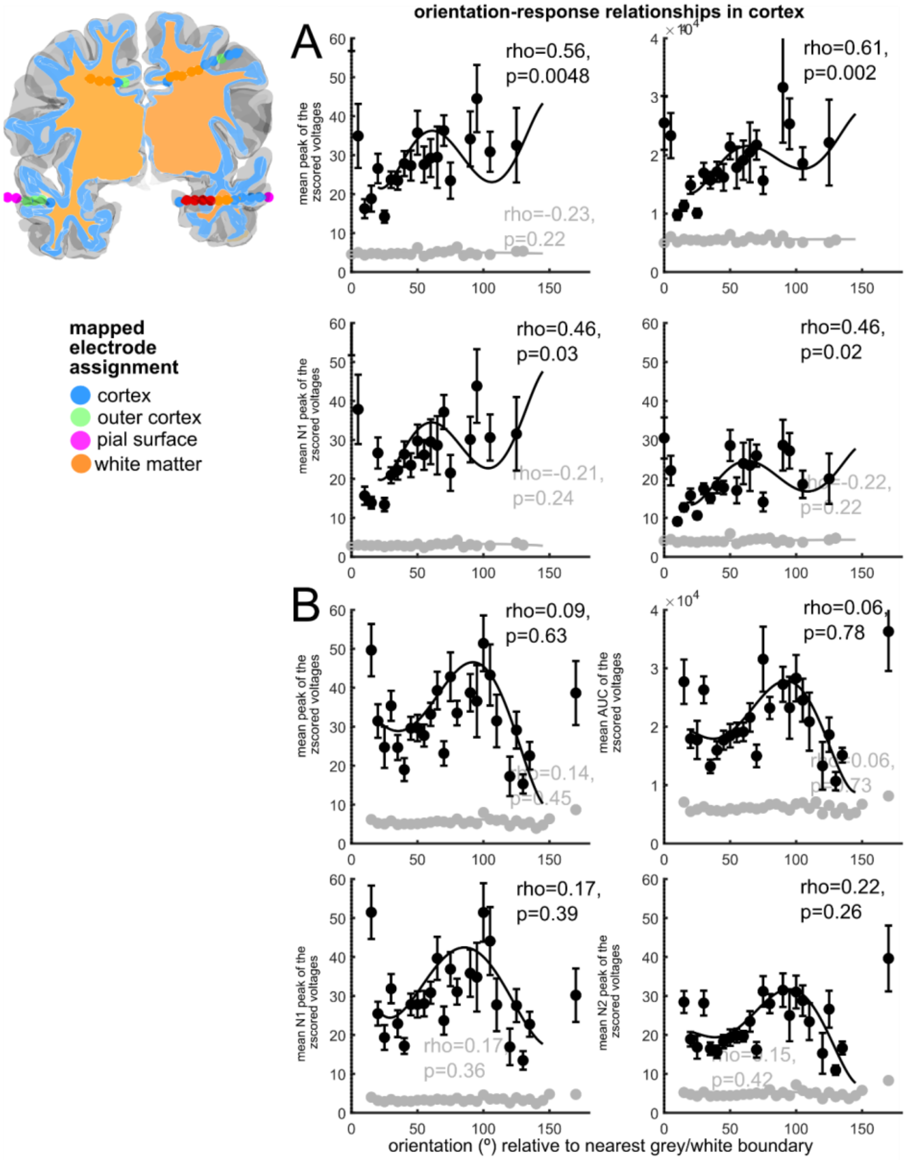


**Supplemental Figure 13. Distances and orientations of stimulating electrodes relative to major landmarks. A**. Location of two example bipolar pairs of electrodes relative to the grey and white matter in a single scanned slice, with an enlarged view of the different measurements which include the closest grey matter and white matter points from the center of the stimulating bipolar pair. **C**. Distribution of distances to the pial surface, the grey/white boundary, and to subcortical structures for all stimulation sites. **D**. Distribution of distances relative to the grey/white boundary across the data set. **E**. Orientation measure relative to the orientation of the closest cortical column and distribution of angles (º) across the data set and the distribution of orientations across all stimulation sites.
